# An inducible genome editing system for plants

**DOI:** 10.1101/779140

**Authors:** Xin Wang, Lingling Ye, Robertas Ursache, Ari Pekka Mähönen

## Abstract

Conditional manipulation of gene expression is a key approach to investigating the primary function of a gene in a biological process. While conditional and cell-type specific overexpression systems exist for plants, there are currently no systems available to disable a gene completely and conditionally. Here, we present a novel tool with which target genes can be efficiently conditionally knocked out at any developmental stage. The target gene is manipulated using the CRISPR-Cas9 genome editing technology, and conditionality is achieved with the well-established estrogen-inducible XVE system. Target genes can also be knocked-out in a cell-type specific manner. Our tool is easy to construct and will be particularly useful for studying genes which have null-alleles that are non-viable or show strong developmental defects.

## MAIN TEXT

Studies of gene function typically rely on phenotypic analysis of loss-of-function mutants. However, mutations may lead to gametophytic or embryonic lethality, or early developmental defects, impeding studies in postembryonic plants. The genome of the model species *Arabidopsis* contains a substantial number of such essential genes, though the precise number remains unknown^1^. Developing a tool that enables conditional and cell-type specific gene disruption is therefore of great value for comprehensively investigating gene function in specific developmental or physiological processes.

Different strategies have been pursued for this purpose. One widely applied approach is the inducible expression of silencing small RNAs^2,3^. However, this results in only a partial reduction of transcript levels, which may hinder a full investigation of gene function. Furthermore, since small RNAs can be mobile^4^, constraining the knockdown effect to a given cell-type is challenging. These limitations can be overcome by using the Cre/lox based clonal deletion system, which provides the possibility of a full knockout together with cell-type specificity. However, this method relies on complicated genetic engineering and has thus remained a rather marginal technique^5,6,7^.

The CRISPR-Cas9 system consists of components derived from the prokaryote adaptive immune system which have been modified for use as a genome editing toolkit in eukaryotes. The endonuclease activity of Cas9 produces double-strand breaks (DSB) in DNA when directed to a target by a single guide RNA (sgRNA). The subsequent error-prone DSB repair mediated by non-homologous end joining facilitates knockout generation. Thus far, CRISPR-Cas9 has been used in plants to generate stable knockouts^8^ and somatic knockouts at fixed developmental stages by driving Cas9 expression with tissue-specific promoters^9^. By integrating the well-established CRISPR-Cas9 technology^10^ with an XVE-based cell-type specific inducible system^11,12^, we developed an Inducible Genome Editing (IGE) system in *Arabidopsis* which enables efficient generation of target gene knockouts in desired cell types and at desired times.

To achieve this, we first generated a fusion of a small nucleolar RNA promoter and an sgRNA (*pAtU3/6-sgRNA*) in two sequential PCR amplification steps (Fig. 1a). The fusion was then cloned into the *p2PR3-Bsa I-ccdB-Bsa I* entry vector (3^rd^ box) by Golden Gate cloning^10^. This method allows simultaneous cloning of several *pAtU3/6-sgRNA* fragments, if needed. Next, we recombined a plant-codon optimized *Cas9p*^10^ into *pDONR 221z* (2^nd^ box). Finally, the IGE binary vector was generated in a single MultiSite Gateway LR reaction by combining an estrogen-inducible promoter (1^st^ box)^11^, *Cas9p* (2^nd^ box), *pAtU3/6-sgRNA* (3^rd^ box) and a plant-compatible destination vector^11,13^ (Fig. 1a). To facilitate screening of transformed seeds, we also generated two non-destructive fluorescent screening vectors (Supplementary Fig. 1). The availability of a large collection of cell-type specific or ubiquitous inducible promoters^11^ and of destination vectors with different selection markers^11,13^ makes the IGE system quite versatile. In summary, an IGE construct can be generated in two cloning steps: first, generating a *pAtU3/6-sgRNA* entry vector by Golden Gate cloning and then performing an LR reaction.

**Figure 1:**
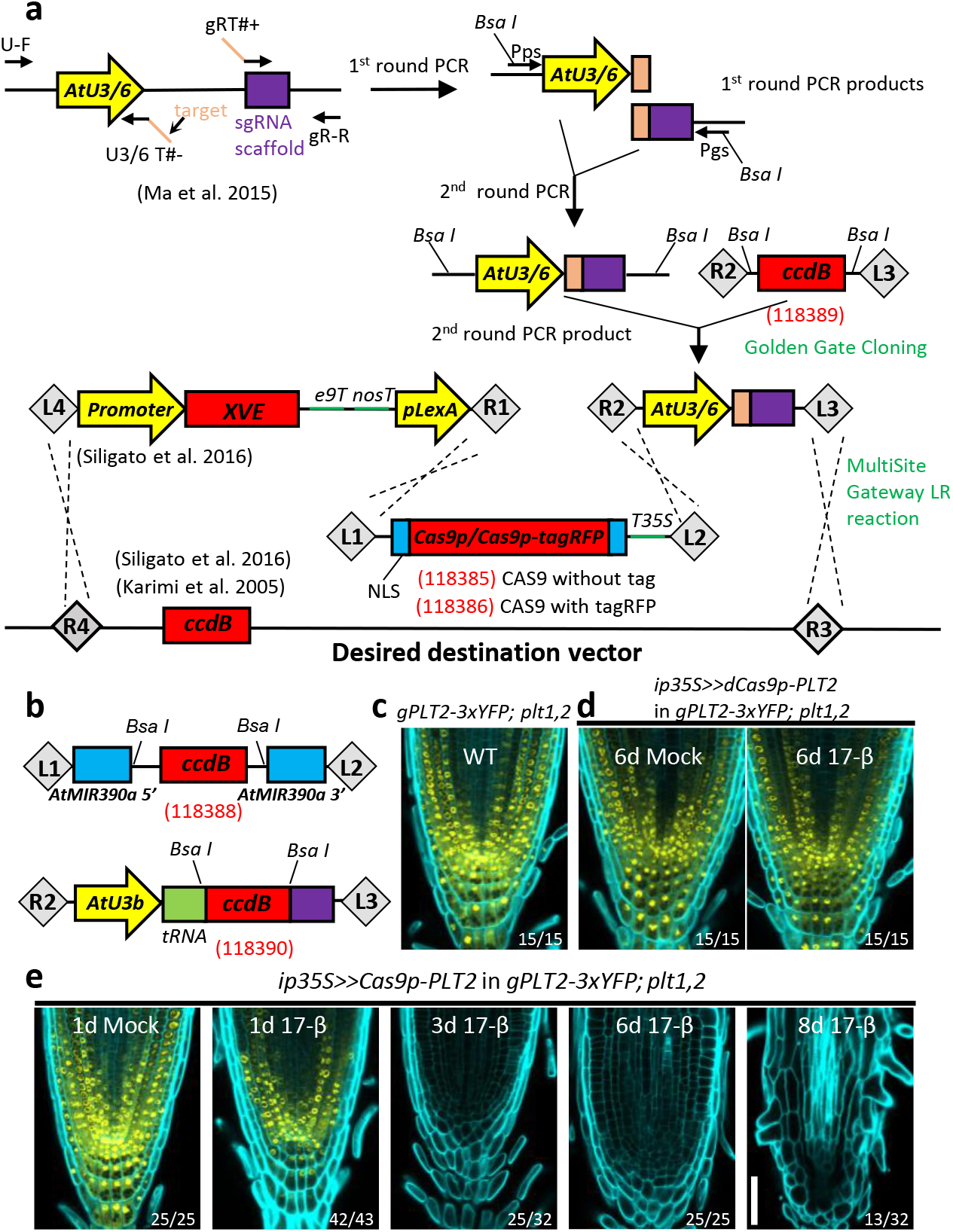
Engineering the IGE system for conditional genome editing. **a**, Cloning steps for IGE construct generation. Fusions of the sgRNA expression cassette (*pAtU3/6-sgRNA*) were constructed by two PCR steps and were subsequently cloned into the *p2R3z-Bsa I-ccdB-Bsa I* entry vector by Golden Gate cloning. The binary IGE construct was then recombined by a MultiSite Gateway LR reaction. **b**, Schematics of two other entry vectors generated in this study. Entry vector *p221z-AtMIR390a*, in which *AtMIR390a* is split by a *Bsa I-*flanking-*ccdB* cassette, was utilized for inducible gene knockdown. Entry vector *p2R3z-AtU3b-tRNA-ccdB-gRNA* was generated to exploit the endogenous tRNA processing system. Two annealed overlapping target sequences with overhangs can be directly ligated into *Bsa I*-linearized *p2R3z-AtU3b-tRNA-ccdB-gRNA*. Red numbers in brackets are the Addgene numbers of vectors created in this study. **c**, The YFP signal in the RM of 7 day-old *gPLT2-3xYFP*;*plt1,2*. **d**, dCas9p does not decrease PLT2-3xYFP expression. **e**, Cas9p-mediated *PLT2* editing resulted in a gradual loss of YFP and eventually full differentiation of the RM. The numbers are the frequency of the observed phenotypes in independent T1 samples. Cell walls are visualized by calcofluor. Scale bar, 50 μm.

Next, we tested the IGE system in the *Arabidopsis* root meristem (RM) by targeting well-established regulatory genes that are essential for RM development. In the RM, a subset of AP2/EREBP family transcription factors, including *PLETHORA1* (*PLT1*) and *PLT2*, form gradients with maxima at the quiescent center (QC) to drive the transition from stem cells to differentiated cells^14–16^. The double mutant *plt1,2* exhibits a fully differentiated RM 6-8 days after germination^14^, which can be rescued by complementing it with *gPLT2-3xYFP*^16^. The fused 3xYFP restricts the mobility of PLT2^16^, making it possible to observe cell-specific effects of editing *PLT2.* We designed four sgRNAs to target *PLT2* in the *gPLT2-3xYFP; plt1,2* background (Supplementary Fig. 2a,b). *Cas9p* or nuclease-dead *Cas9p* (*dCas9p)* were transcribed under the inducible, ubiquitous promoter *35S-XVE* (*ip35S*)^11^. While induction of *dCas9p* had no effect on PLT2-3xYFP levels (Fig. 1d), *Cas9p* induction led to a weakening of the YFP signal almost in every transformant (Fig. 1e). YFP fluorescence was initially reduced in the root cap and occasionally in the epidermis or stele. Prolonged induction gradually abolished the YFP signal and led to RM differentiation after 8-10 days of induction (Fig. 1e and Supplementary Table 1), similar to the uncomplemented *plt1,2* mutant^14^.

**Figure 2:**
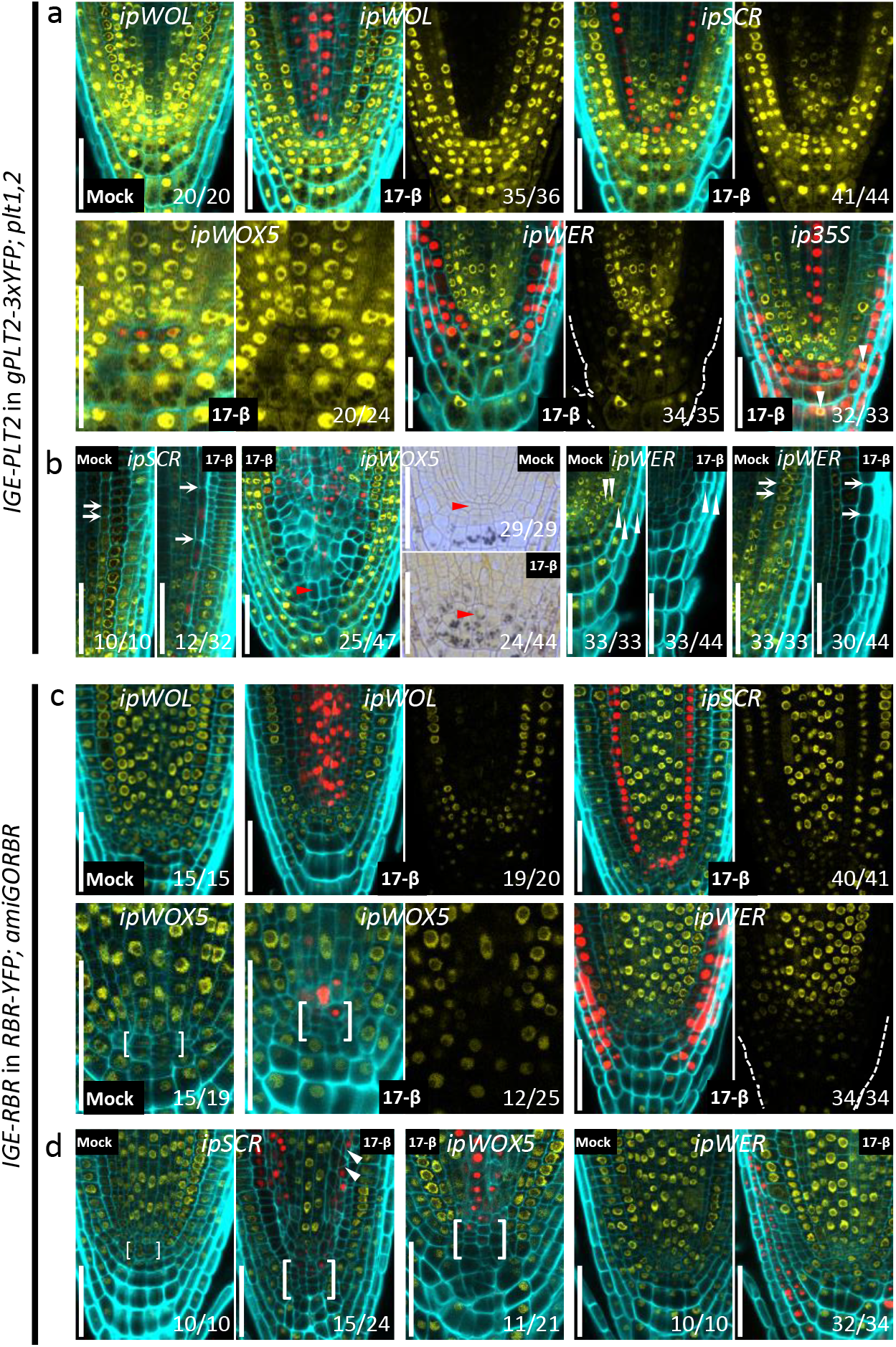
The IGE system enables efficient cell-type-specific genome editing. **a**, A one-day induction is sufficient to induce efficient cell-type specific *PLT2* editing. In rare occasions, we observed overlapping Cas9p-tagRFP and PLT2-3xYFP expression (white arrowhead). **b**, PLT2 is cell-autonomously required for QC and stem cell maintenance. QC cells (red arrowheads) as well as endodermal and epidermal cells (white arrows) showed premature differentiation or cell expansion after 3 days of induction. QC differentiation is accompanied by shift of *ipWOX5* expression towards the provascular cells. Removal of PLT2 from the *ipWER* expression domain also resulted in fewer LRC layers (white arrowhead) and ectopically decreased the PLT2-3xYFP expression. Cas9p-tagRFP expression in the LRC and epidermis was frequently undetectable. **c**, A one-day induction is sufficient to induce efficient cell-type specific *RBR* editing. Without induction, the QC frequently shows cell divisions, probably due to the heterogeneity of the complementing RBR-YFP. **d**, RBR cell-autonomously prevents QC and stem cell division. The endodermis, QC and LRC exhibited overproliferation after 3 days of induction. White arrowheads indicate rotated cell division planes in the endodermis. QC regions are marked by brackets in **c** and **d**. Cell walls are highlighted by calcofluor. The numbers represent the frequency of the observed phenotypes in independent T1 samples. Scale bars, 50 μm.

The requirement of Cas9p nuclease activity for the disappearance of YFP fluorescence suggests that genome editing of *PLT2* caused a homozygous frame-shift mutation in *gPLT2-3xYFP*. PCR was performed to test whether a DNA fragment was deleted within the four target sites in *PLT2*. Intriguingly, only a single truncated band was detected aside from the expected WT band, corresponding to fragment deletion between the first and last targets. Sanger sequencing confirmed this deletion (Supplementary Fig. 2b-d). Further experiments revealed that constructs with just a single sgRNA could achieve equal efficiency in editing *gPLT2-3xYFP*; however, the efficiency strongly depended on the SnoRNA promoter used, with *AtU3b* and *AtU6-29* being the most efficient promoters, at least in the *Arabidopsis* RM (Supplementary Fig. 3 and Supplementary Table 1). Since *AtU3b* and *AtU6-29* were used to drive the expression of sgRNA1 and sgRNA4, respectively, corresponding to the first and last targets in *PLT2*, this seems to explain the prevalence a deletion between these two positions (Supplementary Fig. 2). In summary, the IGE system enables efficient conditional genome editing even with a single sgRNA; however, if a complementary reporter line for the target gene does not exist, it is advisable to use two or more sgRNAs to generate a deletion which can be easily detected by PCR.

Next, we investigated whether the IGE system can be used to induce genome editing in a cell-type specific manner. We tested four inducible promoters: *pWOL-XVE* (*ipWOL*), *pWOX5-XVE* (*ipWOX5*), *pSCR-XVE* (*ipSCR*), and *pWER-XVE* (*ipWER*)^11^, the expression of which, together, covers most of the cell types in the RM. *Cas9p-tagRFP* was used to monitor promoter activity. Constructs were transformed into *gPLT2-3xYFP; plt1,2*. Along with promoter-specific Cas9-tagRFP expression, we observed a corresponding dampening of the YFP signal in the respective domains after one day of induction (Fig. 2a). Consistent with the role of *PLT2* in promoting stem cell maintenance and QC specification, inducible editing in promoter-specific tissues caused premature cell expansion or differentiation of the endodermis, QC, or epidermis/lateral root cap (LRC) after 3 days of induction (Fig. 2b). This reflects the cell-autonomous function of *PLT2* in maintaining an undifferentiated cell state. In addition to QC differentiation, we observed a shift in *ipWOX5* promoter activity towards the provasculature, which resulted in a larger area lacking the YFP signal (Fig. 2b; left panel in *ipWOX5*). The QC and adjacent provascular cells gained columella cell identity, as revealed by the accumulation of starch granules (Fig. 2b; right panels in *ipWOX5*). These results indicate that new QC cells were re-specified from provascular cells following differentiation of the original QC, and the consequent re-specification and differentiation of the QC gradually led to a larger domain without YFP. These results are consistent with experiments in which laser ablation of the QC leads to re-specification of a new QC from provascular cells^17^.

We found that genome editing correlates strongly with Cas9p expression (Supplementary Fig. 4, 5). The expression level, the timing of induction and the expression region of Cas9-tagRFP determined editing performance in independent transformants. In addition, analysis showed that the editing capability of the IGE system is stably transmitted to the T2 generation (Supplementary Fig. 6).

To test whether the IGE system can edit other loci, we targeted a key gene encoding a cell cycle regulator, *RETINOBLASTOMA-RELATED* (*RBR*)^7,18^. The *RBR* null allele is gametophyte-lethal^18^. Previous conditional knockdowns and clonal deletion experiments have shown that RBR has a role in restricting stem cell division in the RM^6,7,19^. RBR-IGE constructs were transformed into a background in which *RBR-YFP* complements an *RBR* artificial microRNA line, *35S:amiGORBR* (*amiGORBR*)^19^. After one day of induction, we observed loss of YFP specifically in the respective promoter domains (Fig. 2c). Three days of induction led to cell overproliferation in the QC, LRC and endodermis, recapitulating the reported phenotype^6,7,19^ (Fig. 2d).

When inducing Cas9p-tagRFP, we found that *ip35S* was not expressed ubiquitously but instead preferentially in the root cap and sometimes in the epidermis or stele (Fig. 2a and Supplementary Fig. 4). This pattern matches the domain of reduced RBR-YFP (Supplementary Fig. 7b) and PLT2-3xYFP expression (Fig. 1e and Supplementary Fig. 4) after a 1-day induction. After long-term induction of *ip35S* or *ipWER*, PLT2-3xYFP expression decreased outside the promoter-active region, in contrast to the effect on RBR (Fig. 1e, Fig. 2b, 2d and Supplementary Fig. 7c). These results suggest that loss of *PLT2* in the epidermis and LRC leads to endogenous, non-cell-autonomous, negative feedback regulation of *PLT2* expression in the rest of the RM, leading to differentiation. In addition, our results confirm the reported cell-autonomous function of RBR^6^.

To further demonstrate the wide applicability of the IGE system, we selected *GNOM* as a target. *GNOM* encodes a brefeldin A (BFA) sensitive ARF guanine-nucleotide exchange factor (ARF-GEF) that plays essential roles in endosomal structural integrity and trafficking^20^. GNOM has been implicated in polar localization of auxin efflux carrier (PINs), but previous studies relied on high-concentration BFA treatments or on hypomorphic alleles^21,22^ because the null allele displays severe overall defects^23,24^. To test the response of PIN1 to the loss of GNOM, we made a construct using the *ipWOL* promoter to target *GNOM* in the vasculature and transformed it into both *GN-GFP*^20^ and *PIN1-GFP*^25^ backgrounds. Following GN-GFP signal disappearance, most transformants displayed short roots, agravitropic growth and reduced lateral root formation 10 days after germination on induction plates (Supplementary Fig. 8, 9), a similar phenotype to the *gnom* mutant^23^. We then focused on PIN1 localization. Following 3 days of induction, PIN1 lost basal polarity and its expression was strongly inhibited (Supplementary Fig. 9), confirming the role of GNOM in driving basal localization of PIN1^21,22^.

When inducing editing of *PLT2*, *RBR* or *GNOM* with *ip35S* or *ipWOL*, we observed cell death in the proximal stem cells of the RM, which have been shown to be sensitive to genotoxic stress^26^ (Supplementary Fig. 10a,b). Although it has been reported that *RBR* silencing causes DNA damage and cell death^27^, *PLT2* and *GNOM* have not been shown to regulate cell death before. It is thus likely that Cas9p-induced DSBs activate downstream DNA damage signals which trigger a cell death response in proximal stem cells.

Next, we tested whether a single YFP-targeting IGE construct can be used to edit several different YFP-containing complementing lines. When targeting fused *YFP* in *gPLT2-3xYFP; plt1,2* and *RBR-YFP*; *amiGORBR* backgrounds, we found a strong reduction in YFP followed by characteristic developmental defects (Supplementary Fig. 11), similar to targeting *PLT2* and *RBR* directly (Fig. 2b, 2d). For example, in *gPLT2-3xYFP; plt1,2*, editing *YFP* in the QC caused QC differentiation, though at a lower frequency than when *PLT2* was targeted. Likewise, we observed LRC overproliferation when targeting *YFP* in *RBR-YFP*; *amiGORBR*. However, unlike when *RBR* was targeted, the YFP signal also decreased in the rest of the RM by an unknown mechanism (Supplementary Fig. 11c). Many fluorescent-tagged lines complementing important genes are available, so targeting reporter-encoding genes might represent a broadly applicable approach for gene function studies. Furthermore, targeting exogenous reporter genes may have fewer off-target effects.

To compare the IGE system with artificial microRNAs (amiRNA) (Fig. 1b), a popular gene knockdown strategy^28,29^, we generated two amiRNAs targeting *PLT2* in *gPLT2-3xYFP*; *plt1,2*. Induction of *amiPLT2-1* by *ip35S* or *ipWOX5* led to a reduction of YFP in a broader domain than with PLT2-IGE (Supplementary Fig. 12a), indicating that IGE is more specific. This is likely due to cell-to-cell movement of amiRNA, consistent with the findings that several microRNAs can move^4^. Additionally, the IGE-caused phenotype tended to be stronger. After a 3-day induction of *ip35S:amiPLT2-1*, the YFP signal was decreased but still visible, and the RM remained undifferentiated after 10 days of induction (Supplementary Fig. 12a and Supplementary Table 1). Likewise, no QC differentiation was observed in *ipWOX5:amiPLT2-1* lines (Supplementary Fig. 12a). The RM of *amiGORBR* showed an overproliferation phenotype, but it was not as severe as in RBR-IGE lines (Supplementary Fig. 12b). To investigate the effect of RBR downregulation in other tissues, we analyzed the root vascular tissue during secondary growth. While *amiGORBR* failed to show any defects in secondary tissue, RBR-IGE caused excessive cell divisions in the phloem and periderm (Supplementary Fig. 12b), indicating a conserved role for RBR in limiting cell divisions in different tissues. Interestingly, the proliferating clones were interspaced with slowly proliferating WT clones, which further confirms the cell-autonomous function of RBR.

In conclusion, we show that the IGE system can be used to disrupt target genes efficiently and precisely. Through spatiotemporal control of Cas9p expression, the system is well-suited to trace early molecular and cellular changes before visible phenotypes appear. Since the estrogen inducible system has been applied in various organs and plant species^12,30,31^, we expect the IGE system to be broadly applicable for plant molecular biology. By using different Cas9 variants, the system can be readily repurposed for base editing or transcriptional regulation.

## METHODS

### Plant material and cloning

To generate the *p221z-Cas9p-t35s* entry vector, first, *Cas9p* with two flanking nuclear localized signal (*NLS*) coding sequence and a *t35* terminator were amplified from vector *pYLCRISRPCas9P35S-B*^10^ with chimeric primers which contained the *attB1/attB2* adaptor at the 5’ end and a 3’ end complementary to *NLS* and *t35s*, respectively. The resultant PCR fragment was gel-purified and then recombined with *pDONR 221* following the instructions of the Gateway BP Clonase II Enzyme mix (Invitrogen).

Site-directed mutations were introduced to two nuclease domains of Cas9p, RuvC1 and HNH (D10A, H840A)^32^, respectively, to generate dCas9. To achieve this, a partial *Cas9p* fragment (61-2582, starting from ATG) was amplified with primers containing the desired mutations. The purified PCR fragment was then used as a mega-primer to amplify *p221z-Cas9p-t35s*. The resulting PCR product was digested by methylation-specific endonuclease Dpn I to remove the parental DNA template before transformation into competent *E.coli* DH5α cells. The presence of mutations in *p221z-dCas9p-t35s* (Addgene ID: 118387) was verified by Sanger sequencing.

To insert the *tagRFP* sequence between *Cas9p* and the 3’ end of the *NLS* encoding sequence located in *p221z-Cas9p-t35s*, *tagRFP* was first amplified from the entry vector *p2R3a-tagRFP-OcsT*^11^ with chimeric primers consisting of a 3’ end of *tagRFP*-specific oligonucleotides and a 5’ end of *Cas9p*/*NLS*-specific oligonucleotides complementary to the flanking sequence at the insertion point. The purified PCR fragment was then used as mega-primer in the subsequent Omega PCR step^33^, which used *p221z-Cas9p-t35s* as the template. The PCR product was treated with Dpn I before transformation into competent *E.coli* DH5α cells. The insertion of *tagRFP* was verified by both enzyme digestion and Sanger sequencing.

To facilitate ligation of the sgRNA expression cassette (*pAtU3/6-sgRNA*) into a Gateway entry vector, the negative selection marker, a *ccdB* expression cassette flanked by two *Bsa I* sites, was amplified from *pYLCRISPRCas9P35S-B*^10^ with primers containing *attB2/attB3* adaptors. After a BP reaction with *pDONR P2R-P3z*, the reaction mixture was transformed into the ccdB-tolerant *E.coli* strain DB3.1. Colony PCR was performed to screen for positive colonies which had been transformed with recombined plasmids but not the empty *pDONR-P2R-P3z*. The presence of the *p2R3z-Bsa I-ccdB-Bsa I* entry vector was then further confirmed by enzyme digestion and Sanger sequencing.

The sgRNA expression cassettes were obtained as previously described^10^. Briefly, the first round of PCR amplified *AtU3/6* promoters from template vectors, *pYLsgRNA-AtU3b* (Addgene ID: 66198), *pYLsgRNA-AtU3d* (Addgene ID: 66200), *pYLsgRNA-AtU6-1* (Addgene ID: 66202) or *pYLsgRNA-AtU6-29* (Addgene ID: 66203), using a common forward primer, *U-F*, and reverse chimeric primer *U3/6 T#-* which contains an *AtU3/6-*specific sequence at the 3’ end and a target sequence at the 5’ end. All sgRNA scaffolds were amplified from *pYLsgRNA-AtU3b* with a common reverse primer, *gR-R*, and chimeric forward primer *gRT #+*, which includes the sgRNA specific sequence at the 3’ end and the target sequence at the 5’ end. In the second round of PCR, purified first-round PCR products were used as templates for overlapping PCR with Bsa I-containing primers *Pps/Pgs* as primer pairs. In this study, four sgRNAs (sgRNA1-sgRNA4) transcribed under promoters *AtU3b*, *AtU3d*, *AtU6-1*, and *AtU6-29*, respectively, were used to target genes of interest. For each target gene, four relatively equally distributed target sites were manually selected by following rules described previously^10^. Different sgRNA expression cassettes were cloned into the *p2R3z-Bsa I-ccdB-Bsa I* entry vector by one-step Golden Gate cloning. Golden gate cloning was performed with 120ng *p2R3z-Bsa I-ccdB-Bsa I*, 90 ng purified PCR product of each sgRNA expression cassette, 1.5µl 10x fast digestion buffer of Bsa I, 1.5µl Bsa I enzyme (15U), 1.5µl 10mM ATP, 4µl T4 DNA ligase (20U), and H_2_O to make up 15µl. The reaction mixture was incubated at 37 °C for 4-6h before *E. coli* transformation. Selection of positive transformants was performed as described above.

To generate the *p221z-AtMIR390a* entry vector (Fig. 1b), a BP reaction was performed with *pDONR 221* and *pMDC123SB-AtMIR390a-B*/*c*^28^ (Addgene ID: 51775). *pMDC123SB-AtMIR390a-B*/*c* contains *AtMIR390a* 5’ end and *AtMIR390a* 3’ end which were split by *Bsa I*-flanking *ccdB* expression modules. After transforming DB3.1, positive colonies were screened by colony PCR followed by enzyme digestion and sequencing. Two artificial microRNA against *PLT2* (*amiPLT2*-*1* and *amiPLT2*-*2*) were designed using http://p-sams.carringtonlab.org/. Annealed *amiPLT2* was ligated into *p221z-AtMIR390a* by a one-step reaction as previously described^28^.

Tandem arrayed tRNA-sgRNA units have been exploited for multiplex genome editing by using the endogenous tRNA processing machinery^34^, which precisely cuts tRNA precursors at both ends and releases free sgRNA after transcription. This strategy has been applied in a variety of plant species^34,35^. However, to date there are few reports of its application in *Arabidopsis*. We therefore investigated its feasibility in *Arabidopsis* genome editing and meanwhile tested its compatibility with our IGE system. To facilitate target sequence ligation, we first constructed a *p2R3z*-*AtU3b-tRNA-ccdB-sgRNA* entry vector (Fig. 1b). AtU3b, tRNA-1, tRNA-2 (tRNA was amplified in two separate fragments), the ccdB expression cassette (flanked by Bsa I), and the sgRNA scaffold were amplified with the indicated primer pairs. Both ends of each fragment contained primer-introduced sequences overlapping with the desired flanking fragments. In the overlapping PCR step, *attB2-AtU3b-F* and *attB3-sgRNA-R* were used as a primer pair to assemble these five purified PCR fragments, which were mixed as templates. Cloning this fused fragment into *pDONR P2R-P3z* was conducted as described above. To clone the first target sequence of *PLT2* into *p2R3z*-*AtU3b-tRNA-ccdB-sgRNA*, two annealed primers with 4-nucleotide overhangs at the 5’ ends and 20-nucleotide complementary target sequences were ligated into the entry vector in a one-step reaction as described previously^28^. In the *Arabidopsis* RM, we observed a decrease of the YFP signal in the region where the inducible promoter was active in most independent lines after a 1-day induction and finally a fully differentiated RM after a 10-day induction (Supplementary Fig. 3; Supplementary table 1), indicating that sgRNA against *PLT2* was disassociated from tRNA processing and guiding Cas9p to cleave *PLT2*. It has recently been reported that efficient genome editing could be achieved by fusing tRNA to a mutant sgRNA scaffold but not the wild type sgRNA scaffold in *Arabidopsis*^36^. However, in our hands wild type sgRNA scaffold and tRNA fusion worked well. We reasoned that the sgRNA promoter, Cas9 variant, sgRNA scaffold, target loci, and the tissue to be edited may all affect tRNA-sgRNA-mediated editing performance in *Arabidopsis*. Therefore a future comprehensive study of these variables may improve the utility of the tRNA processing system in *Arabidopsis*.

The red seed coat vector *pFRm43GW* (Addgene ID: 133748) was generated by modifying the *pHm43GW* destination vector^13^, which was obtained from VIB (https://gateway.psb.ugent.be/). The *pHm43GW* vector was digested with PaeI (SphI) (ThermoFisher Scientific) to remove the hygromycin cassette. Using an In-Fusion HD Cloning (TaKaRa) kit, two fragments were cloned into the digested vector. The first fragment contained a *ccdB* cassette and recombination sites for MultiSite Gateway cloning, and it was amplified from *pHm43GW* using GAACCCTGTGGTTGGCATGCACATACAAATGGACGAACGGATAAA as a forward primer and ATACCTACATACACTTGAAGGGTACCCGGGGATCCTCTAGAGGG as a reverse primer. The second fragment contained the FastRed module, consisting of the *OLE1* promoter followed by *OLE1 tagRFP*, which was amplified from *pFAST-R01*^37^ using CTTCAAGTGTATGTAGGTATAGTAACATG as a forward primer and CGAATTGAATTATCAGCTTGCATGCAGGGTACCATCGTTCAAACATTTGGCAAT as a reverse primer.

We also provide another non-destructive fluorescent screening vector, the green seed coat vector *pFG7m34GW* (Addgene ID: 133747). It was generated by cloning the FastGreen module into the *pP7m34GW* vector^13^, which was obtained from VIB (https://gateway.psb.ugent.be/). The *pP7m34GW* vector was digested with SacI (ThermoFisher Scientific). Three fragments were cloned into the digested *pP7m34GW*. The first fragment contained the *OLE1* promoter followed by the *OLE1* genomic sequence and was amplified from *pFRm43GW* using CCATATGGGAGAGCTCCTTCAAGTGTATGTAGGTATAGT as a forward primer and GCCCTTGCTCACCATAGTAGTGTGCTGGCCACCACGAG as a reverse primer; the second fragment contained the *EGFP* encoding sequence and was amplified from the *pBGWFS7* vector^13^ using ATGGTGAGCAAGGGCGAGGAGCTGT as a forward primer and ATCTATGTTACTAGATCACTTGTACAGCTCGTCCATGCC as a reverse primer; the third fragment contained the *nosT* terminator sequence and was amplified from the *p1R4-ML:XVE* vector^11^ using TCTAGTAACATAGATGACACCGCGCG as a forward primer and TTAACGCCGAATTGAATTCGAGCTCCATCGTTCAAACAT as a reverse primer. All three fragments were combined together with the digested vector using In-Fusion HD Cloning.

The five inducible promoters (*p1R4-p35S:XVE*, *p1R4-pSCR:XVE*, *p1R4-pWER:XVE*, *p1R4-pWOL:XVE*) were created earlier^11^. To construct the binary vector, a MultiSite Gateway LR reaction was performed with the inducible promoters in the 1^st^ box, *Cas9p*, *dCas9p*, *Cas9p-tagRFP* or *amiPLT2* in the 2^nd^ box, the sgRNA expression cassette or *nosT* terminator in the 3^rd^ box and *pBm43GW* (PPT (phosphinotricin) selection) or *pFRm43GW* (seed coat RFP selection) as the destination vectors. All constructs generated in this study are listed in Supplementary table 3.

*PLT2*-targeting constructs were dipped into the *gPLT2:3xYFP,plt1,2* background^16^. For *RBR*-targeting constructs, the dipping background was segregating *pRBR:RBR-YFP*(+,-); *35S:amiGORBR*(+,+)^19^. The IGE construct targeting *GNOM* was transformed into both the *GN-GFP*^20^ and *PIN1-GFP*^25^ backgrounds. With the exception of the construct transformed into the GN-GFP background, in which the GFP signal was weak, all T1 lines were prescreened under a fluorescence-binocular microscope to identify those with leaky inducible promoter or in which the root tip had been damaged during selection. Only lines with YFP/GFP signal in root tip were used for further experiments. The *PLT2* and *RBR*-based backgrounds were also used for *YFP*-targeting construct transformation. The *RBR*-targeting construct *ip35S>>Cas9p-RBR* was also dipped into the *Col-0* background. All experiments were conducted using T1 plants unless stated otherwise. Each experiment has been repeated at least three times, except the RM differentiation characterization in Supplementary Table 1, which was repeated twice.

### Plant growth and chemical treatments

All seeds were surface-sterilized with 20% chlorine for 1 min, followed by a 1 min incubation in 70% ethanol and two rinses in H_2_O. The sterilized seeds were kept at 4°C for two days before plating on half strength Murashige and Skoog growth medium (½ GM) plates with/without selection antibiotics. The plates were vertically positioned in a growth chamber at 22 °C in long day conditions. PPT selection was conducted by growing sterilized seeds on ½ GM plates containing 20 μg/ml PPT for 4 days, then transferring them to PPT-free ½ GM plates for another 2 days before treatment. The trans-pFRm43GW-based seeds were screened under a fluorescence binocular using DSRed filter (Supplementary Fig. 1b), and the sterilized seeds were directly grown on ½ GM plates for 6 days before treatment. 17-β-estradiol (17-β, Sigma) was dissolved in dimethyl sulfoxide (DMSO, Sigma) to make 10 mM stock solution (stored at −20°C) and a 5 μM working concentration was used. An equal volume of DMSO was used as a mock treatment.

### Microtome sectioning and histological staining

Transverse plastic sections were cut from *ip35S>>Cas9p-RBR* (in *Col-0* background) roots which were geminated on estradiol plates for 20 days, as well as *Col-0* and *35S:amiGORBR* roots that were grown on ½ GM plates for 20 days. Sections from 5 mm below the root–hypocotyl junction point were used for analysis. Sections were stained in 0.05% (w/v) ruthenium red solution (Fluka Biochemika) for 5 seconds before microscopy analysis. For root samples from *ipWOX5>>Cas9p-tagRFP-PLT2*, *ipWOX5>>Cas9p-tagRFP-YFP* and *ipWOX5>>amiPLT2-1*, after 3 days of mock or 17-β treatment, a serial longitudinal section of 5 µm thickness was cut from the root tips. To observe the QC differentiation state, the longitudinal sections were stained in 1g/ml lugol solution (Sigma) for 12 seconds before observation under a microscope. The sectioning methodology has been previously described^38^.

### Microscopy and image processing

All of the cross sections and longitudinal sections were visualized using a Leica 2500 microscope. All fluorescent images were taken with a Leica TCS SP5 II Confocal microscope. Root samples used for cell death detection were stained in 10 μg/mL propidium iodide for 10 mins then rinsed twice in water before imaging. For other samples used for fluorescence observation, a ClearSee protocol^39^ was used with slight modifications. Roots were first fixed in 4% paraformaldehyde (dissolved in 1xPBS, PH 7.2) for at least one hour with vacuuming, then washed twice in 1x PBS and transferred to ClearSee solution. Samples were incubated in ClearSee solution for at least 24h. Before imaging, 0.1% calcofluor white dissolved in ClearSee was used for one hour with vacuuming to stain cell walls. This was followed by washing the samples in ClearSee solution for at least 30 mins with shaking. During the washing, the ClearSee solution was changed every 15 mins. Confocal settings were kept the same between mock and induction in each experiment. All confocal images were acquired in sequential scanning mode. Images were sometimes rotated using Photoshop and the resulting empty corners were filled with a black background. All images were cropped and organized in Microsoft PowerPoint. The brightness of the calcofluor signal was sometimes adjusted differently between the mock and induction for better cell wall visualization.

## ACKNOWLEDGEMENT

We thank B. Scheres (Wageningen University, the Netherlands) and N. Geldner (University of Lausanne, Switzerland) for sharing published materials; N. Geldner for providing support for R.U.; and S. el-Showk for proofreading of this manuscript. This work was supported by the Academy of Finland (grants #316544, #266431, #307335), University of Helsinki HiLIFE fellowship (X.W., L.Y., A.P.M.) and European Molecular Biology Organisation (EMBO ALTF 1046-2015 to R.U.). X.W. is also supported by a grant from the Chinese Scholarship Council (CSC).

## CONTRIBUTIONS

X.W. and A.P.M. designed the experiments. X.W. conducted all experiments, except L.Y. carried out the analysis for Supplementary Table 1. R.U. generated and tested the new destination vectors. X.W. and A.P.M. analyzed the results and wrote the manuscript, with input from all co-authors.

---

**Supplementary Figure 1 Non-destructive screening markers facilitate identification of transformed seeds.**

**(a)** Non-destructive fluorescent screening destination vectors generated in this study. **(b)** Examples of trans-pFRm43GW seeds screened under the fluorescence-binocular in the T1 (left) and T2 (right) generations.

**Supplementary Figure 2 PCR genotyping of *PLT2* deletions.**

**(a)** Tandem arrayed sgRNA expression cassettes. (**b)** The genomic structure of *PLT2*. Boxes indicate exons. Orange bars represent target sites in *PLT2*. Black arrows represent relative positions of the forward and reverse primers. **(c)** PCR detection of *PLT2* deletion in *ip35S>>Cas9p-PLT2*; *gPLT2-3xYFP*;*plt1,2* T1 seedlings after 3 days of treatment (in 6 day-old plants). Pooled DNA was isolated from 2cm root segments below the hypocotyl of 10 seedlings. Three primer pairs were used. There were no detectable truncated bands in 7-day old *gPLT2 3xYFP*;*plt1,2*, while weak truncated bands were detected in mock treated seedlings (white arrowhead), probably due to weak leakiness of *ip35S* in certain roots or cells. Note that although four sgRNAs were used to target *PLT2*, only one truncated band was detected with each primer pair. **(d)** Sequencing of truncated bands from primer pair F-R3 confirmed deletion between the 1^st^ and 4^th^ *PLT2* target sites (letters in red represent protospacer adjacent motif, PAM). To determine the deletion types, the truncated band was not directly used for sequencing but cloned into *pDONR 221*. Two deletion types were found in 4 sequenced recombinant vectors.

**Supplementary Figure 3 sgRNA promoter identity affects editing efficiency in *Arabidopsis* roots.**

For each construct, the indicated sgRNA promoter was used to drive transcription of sgRNA1, while *ip35S* was used to guide *Cas9p* transcription. *AtU3b* and *AtU6-29* showed the best editing efficiency in T1 seedlings after one day of induction. Transcription of tRNA together with sgRNA1 under the *AtU3b* promoter also resulted in efficient *PLT2* editing. WT is the 7-day old *gPLT2-3xYFP*; *plt1,2*. White dotted lines mark the RM outlines. Cell walls are highlighted by calcofluor. Numbers indicate the frequency of similar results in the independent T1 samples analyzed. Scale bar, 50 μm.

**Supplementary Figure 4 IGE-mediated genome editing correlates with Cas9 expression.**

After one day of induction, IGE performance on *PLT2* editing under different inducible promoters was classified into two categories. In the mild category, Cas9p-tagRFP expression tends to be weak and narrow, resulting in narrow domains of moderately decreased YFP signal. In the strong category, Cas9p-tagRFP expression was strong and broad, with strongly and broadly reduced YFP fluorescence. In the uppermost panel, Cas9p was used without a tag. White dotted lines mark the RM outlines. Cell walls are visualized by calcofluor. Numbers indicate the frequency of similar results in the T1 samples analyzed. Scale bars, 50 μm.

**Supplementary Figure 5 IGE system enables real time observation of genome editing.**

To monitor *PLT2* editing dynamics, a time-course 17-β induction was conducted to *ipWER >> Cas9p-tagRFP-PLT2* in *gPLT2-3xYFP*; *plt1,2* (T2 generation, #1). The reduction of PLT2-3xYFP expression was first detected after 12 hours of induction and became obvious with 16 hours of induction. The editing activity was gradually spread inwards, likely due to the radial diffusion of 17-β within *ipWER* domain. White dotted lines mark the RM outlines. Cell walls are visualized by calcofluor. Numbers indicate the frequency of observed phenotype within given induction duration. Scale bar, 50 μm.

**Supplementary Figure 6 The capacity of conditional genome editing by IGE system is inherited.**

For each construct, two independent transgenic T2 lines were randomly selected and checked. Representative images are shown. Note that the second *ipWOX5>>Cas9p-tagRFP-PLT2* line was leaky: roots displayed a similar phenotype with/without induction. Cell walls are marked by calcofluor. Numbers represent the frequency of the observed phenotype in analyzed T2 samples. Scale bar, 50 μm.

**Supplementary Figure 7 RBR functions cell-autonomously in the RM.**

**(a)** A three-day mock treatment of *ip35S>>Cas9p-RBR* in *RBR-YFP; amiGORBR*. **(b)** A one-day induction caused a reduced RBR-YFP signal mainly in the root cap region without an obvious phenotype. **(c)** Inducing *RBR* editing with *ip35S* typically led to LRC overproliferation (white arrows) without affecting the YFP signal in other domains after a 3-day induction. In some cases, both wild type cells and RBR-knockout cells were seen on the same root (left in **c**). Cell walls are visualized by calcofluor. Numbers indicate the frequency of the observed phenotype in independent T1 samples. Scale bar, 50 μm.

**Supplementary Figure 8 Post-embryonically inducing *GNOM* editing recapitulates the phenotypes of the *gnom* mutant.**

**(a)** Plants with *ipWOL>>Cas9p-tagRFP-GNOM*; *PIN1-GFP* ten days after germination on mock or 17-β plates. Inducing *GNOM* editing led to shorter roots, agravitropic growth and decreased lateral root (LR) numbers. Adventitious roots from the hypocotyl were frequently found, but these roots were not counted in LR quantification. For each independent root, LR number and root length is quantified in **(b)**. Scale bar, 1 cm.

**Supplementary Figure 9 GNOM is required for PIN1 polarity and expression.**

**(a)** *GNOM* expression disappeared from the vasculature after a 6-day induction of IGE targeting *GNOM*. Due to the weak GFP signal, only roots showing a clear loss of GFP signal were included in quantification. **(b)** A three-day induction of *ipWOL>> Cas9p*-*tagRFP-GNOM*; *PIN1-GFP* resulted in loss of polarity and decreased expression of PIN1-GFP in the endodermis (en), pericycle (p) and stele (s) (white arrows). Right panels are magnified images of the regions marked with a red box in the left panels. Cell walls are marked by calcofluor. Numbers indicate the frequency of the observed phenotype in independent T1 samples analyzed. Scale bar in right panels of **a**, 25 μm; others, 50 μm.

**Supplementary Figure 10 Cas9p-mediated genome editing in proximal stem cells induces cell death.**

**(a)** Stem cell death surrounding the QC was observed after one day of *ip35S>>Cas9p-PLT2* induction. Based on cell types, the cell death response is classified into three categories: provascular cell death, LRC/epidermis initial cell death and columella initial cell death. Samples were counted twice if they had cell death in different domains. **(b)** Cell death of provascular cells and early descendants was induced after one day of induction of *ipWOL>>Cas9p-tagRFP-PLT2/RBR/GNOM*. Cell walls are highlighted by propidium iodide (PI). Under PI detection settings, Cas9p-tagRFP is also visible. Numbers indicate the frequency of the observed phenotype in independent T1 samples analyzed. Scale bars, 50 μm.

**Supplementary Figure 11 A single IGE construct targeting a gene encoding a fluorescent reporter has the potential to disrupt different transgene targets.**

**(a)** Editing *YFP* instead of *PLT2* in the *ipWER* expression region caused changes similar to direct *PLT2* editing. The RM had fewer LRC layers (white arrowheads), as well as premature expansion of epidermal cells and a broad, faint YFP signal. The Cas9p-tagRFP signal is frequently invisible. **(b)** Editing *YFP* led to QC (black arrow) differentiation at a lower frequency. **(c)** Targeting the *YFP* of RBR-YFP in the LRC led to LRC overproliferation, similar to editing RBR. However, the YFP signal outside *ipWER* expression region was also hampered by an unknown mechanism, unlike when editing *RBR*. White arrows mark the neighboring cell walls in **a** and **c**. The same construct was used in **a** and **c**. Cell walls are highlighted by calcofluor. Numbers indicate the frequency of the observed phenotype in independent T1 samples analyzed. Scale bars, 50 μm.

**Supplementary Figure 12 Comparison of IGE system with inducible amiRNA.**

**(a)** IGE-PLT2 displays more specific and stronger *PLT2-YFP* downregulation than amiPLT2. After a one-day induction, *ip35S>>amiPLT2-1*; *gPLT2-3xYFP*;*plt1*,*2* and *ipWOX5>>amiPLT2-1*; *gPLT2*-*3xYFP*;*plt1*,*2* showed a broader reduction of the YFP signal, particularly in the bracketed regions where no inducible promoter activity was found. Conversely, induced *PLT2* editing caused very local loss of the YFP signal. After a three-day induction, the YFP signal is still visible in most of *ip35S>>amiPLT2-1*; *gPLT2-3xYFP; plt1,2* transformants but not in *ip35S>>Cas9p-PLT2*; *gPLT2*-*3xYFP*;*plt1*,*2* transformants. There was no QC differentiation in *ipWOX5>>amiPLT2-1*; *gPLT2-3xYFP*; *plt1,2* roots. WT here means 7-day old *gPLT2*-*3xYFP; plt1,2*. White arrows mark the QC. **(b)** Comparison of the RM and root secondary growth of *Col-0*, *35S:amiGORBR* and *ip35S>>Cas9p-RBR*. Inducing *RBR* editing (germination and six days of growth on 17-β plates) resulted in more excessive cell divisions in the LRC than was seen in *amiGORBR* roots (germination and six days of growth on 17-β free plates). Furthermore, RBR editing caused cell overproliferation in secondary tissues such as phloem (ph) cells and the periderm (pe), which was not observed in *amiGORBR* roots. The knockout (ko) sectors (green dotted line) were frequently accompanied by WT sectors (red dotted line), which can be regarded as an internal control. Cell walls are marked by calcofluor. Numbers indicate the frequency of observed phenotype in independent samples analyzed. Scale bars, 50 μm.

**Supplementary Table 1 Quantification of fully differentiated RM after 10 days induction.**

**Supplementary Table 2 Primer list in this study.**

Underlined sequences indicate Gateway adaptors. Sequence in red represent the target sequence in the gene.

**Supplementary Table 3 Constructs list in this study.**

