## Supplementary Figures 1-12, Tables 1-3 for "An inducible genome editing system for plants"

### Supplementary Figure 1

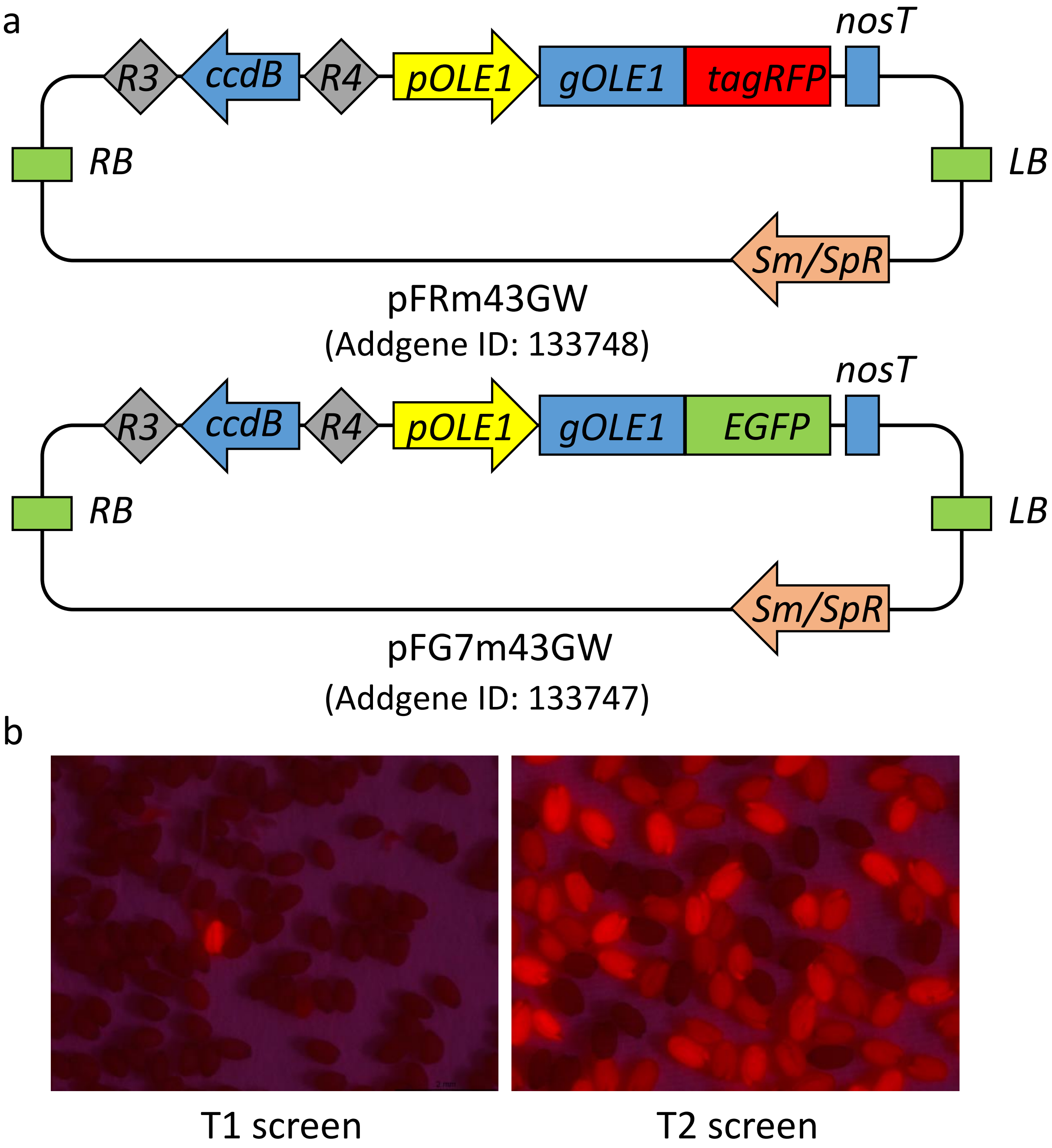


**Supplementary Figure 1 Non-destructive screening markers facilitate identification of transformed seeds.**

**(a)** Non-destructive fluorescent screening destination vectors generated in this study. **(b)** Examples of trans-pFRm43GW seeds screened under the fluorescence-binocular in the T1 (left) and T2 (right) generations.

### Supplementary Figure 2

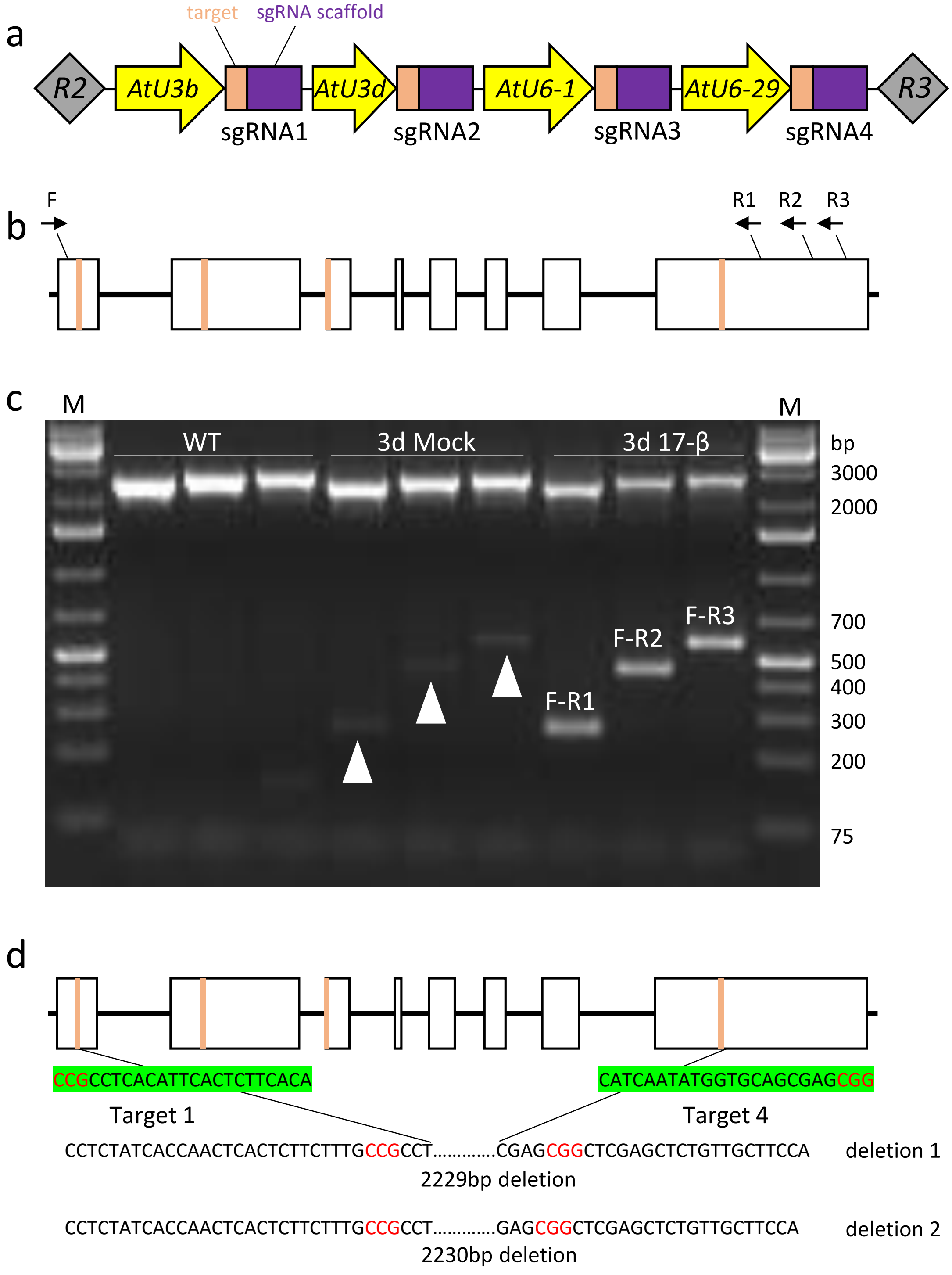


**Supplementary Figure 2 PCR genotyping of *PLT2* deletions.**

**(a)** Tandem arrayed sgRNA expression cassettes. **(b)** The genomic structure of *PLT2*. Boxes indicate exons. Orange bars represent target sites in *PLT2*. Black arrows represent relative positions of the forward and reverse primers. **(c)** PCR detection of *PLT2* deletion in *ip35S>>Cas9p-PLT2*; *gPLT2-3xYFP*;*plt1,2* T1 seedlings after 3 days of treatment (in 6 day-old plants). Pooled DNA was isolated from 2cm root segments below the hypocotyl of 10 seedlings. Three primer pairs were used. There were no detectable truncated bands in 7-day old *gPLT2 3xYFP*;*plt1,2*, while weak truncated bands were detected in mock treated seedlings (white arrowhead), probably due to weak leakiness of *ip35S* in certain roots or cells. Note that although four sgRNAs were used to target *PLT2*, only one truncated band was detected with each primer pair. **(d)** Sequencing of truncated bands from primer pair F-R3 confirmed deletion between the 1st and 4th *PLT2* target sites (letters in red represent protospacer adjacent motif, PAM). To determine the deletion types, the truncated band was not directly used for sequencing but cloned into *pDONR 221*. Two deletion types were found in 4 sequenced recombinant vectors.

### Supplementary Figure 3

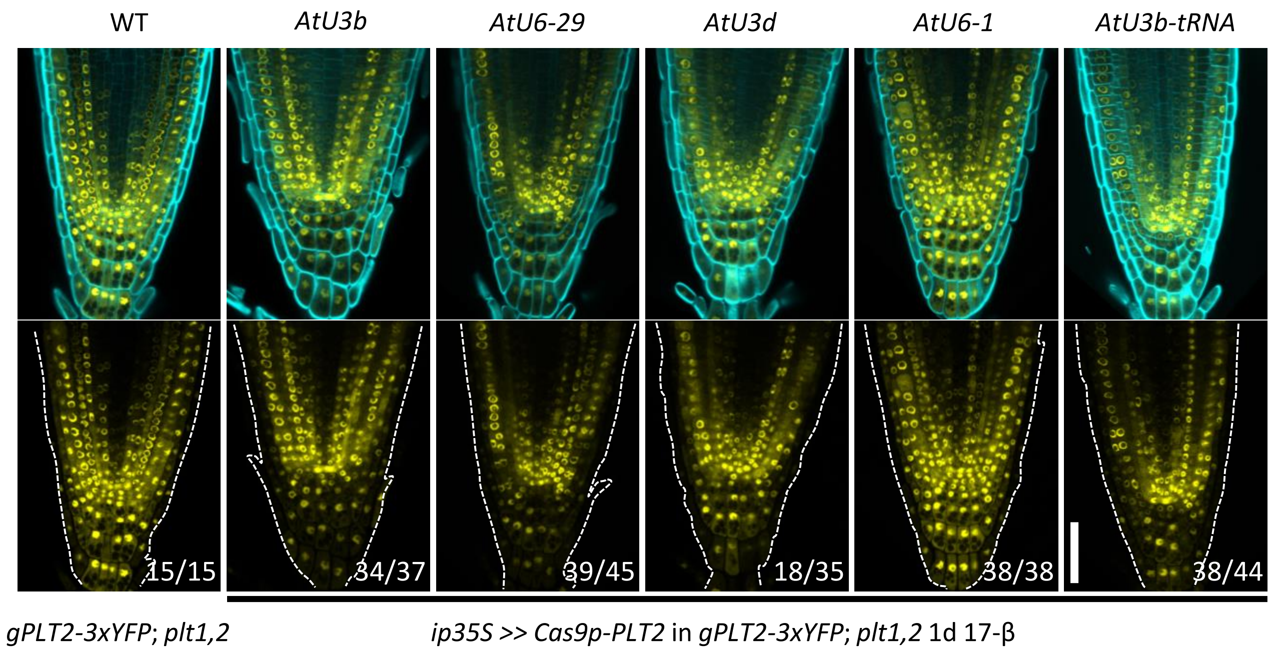


**Supplementary Figure 3 sgRNA promoter identity affects editing efficiency in *Arabidopsis* roots.**

For each construct, the indicated sgRNA promoter was used to drive transcription of sgRNA1, while *ip35S* was used to guide *Cas9p* transcription. *AtU3b* and *AtU6-29* showed the best editing efficiency in T1 seedlings after one day of induction. Transcription of tRNA together with sgRNA1 under the *AtU3b* promoter also resulted in efficient *PLT2* editing. WT is the 7-day old *gPLT2-3xYFP*; *plt1,2*. White dotted lines mark the RM outlines. Cell walls are highlighted by calcofluor. Numbers indicate the frequency of similar results in the independent T1 samples analyzed. Scale bar, 50 μm.

### Supplementary Figure 4

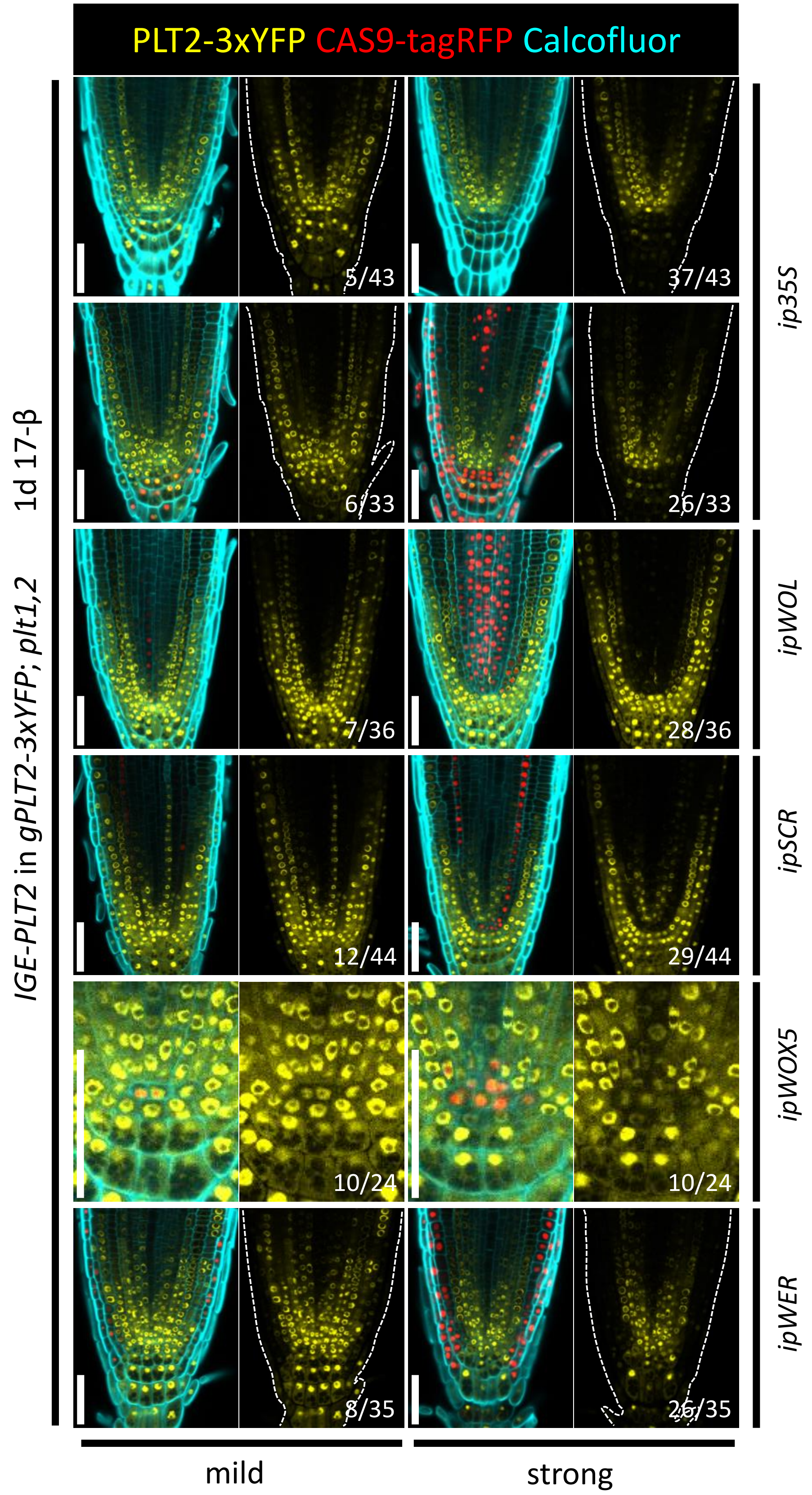


**Supplementary Figure 4 IGE-mediated genome editing correlates with Cas9 expression.**

After one day of induction, IGE performance on *PLT2* editing under different inducible promoters was classified into two categories. In the mild category, Cas9p/Cas9p-tagRFP expression tends to be weak and narrow, resulting in narrow domains of moderately decreased YFP signal. In the strong category, Cas9p-tagRFP expression was strong and broad, with strongly and broadly reduced YFP fluorescence. In the uppermost panel, Cas9p was used without a tag. White dotted lines mark the RM outlines. Cell walls are visualized by calcofluor. Numbers indicate the frequency of similar results in the T1 samples analyzed. Scale bars, 50 µm.

### Supplementary Figure 5

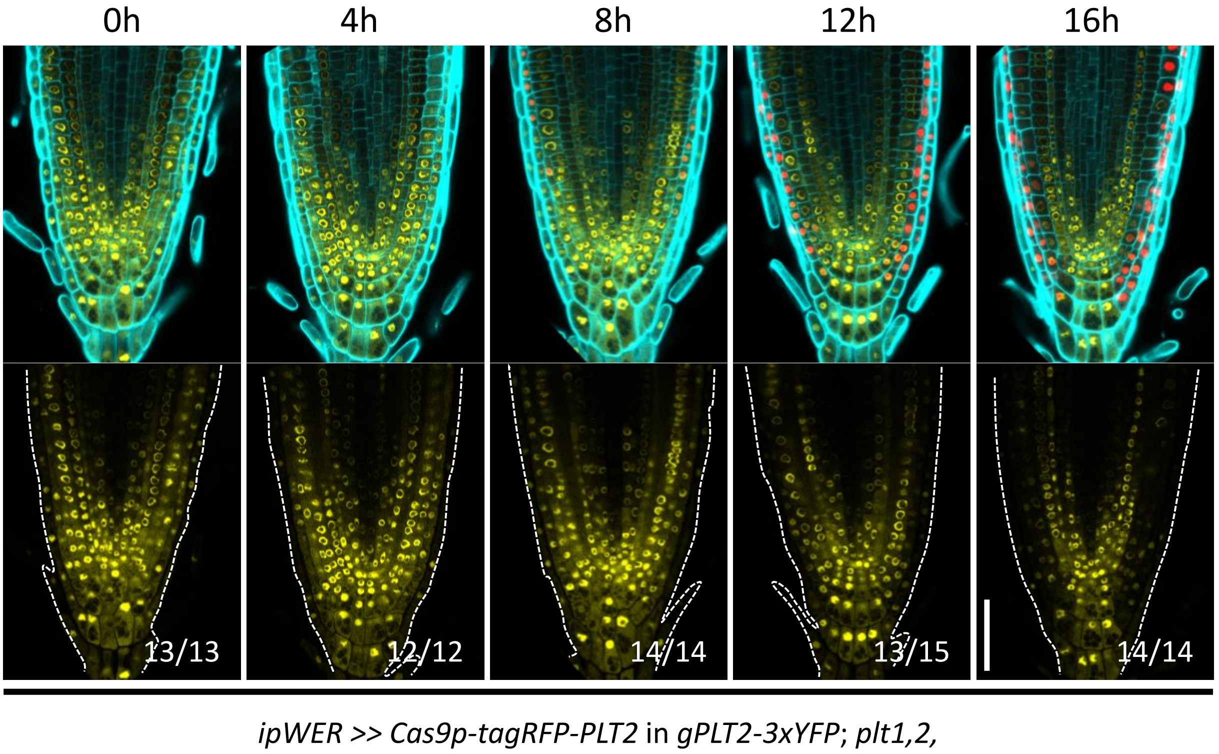


**Supplementary Figure 5 IGE system enables real time observation of genome editing.**

To monitor *PLT2* editing dynamics, a time-course 17-β induction was conducted to *ipWER* >> *Cas9p-tagRFP-PLT2* in *gPLT2-3xYFP*; *plt1,2* (T2 generation, #1). The reduction of PLT2-3xYFP expression was first detected after 12 hours of induction and became obvious with 16 hours of induction. The editing activity was gradually spread inwards, likely due to the radial diffusion of 17-β within *ipWER* domain. White dotted lines mark the RM outlines. Cell walls are visualized by calcofluor. Numbers indicate the frequency of observed phenotype within given induction duration. Scale bar, 50 μm.

### Supplementary Figure 6

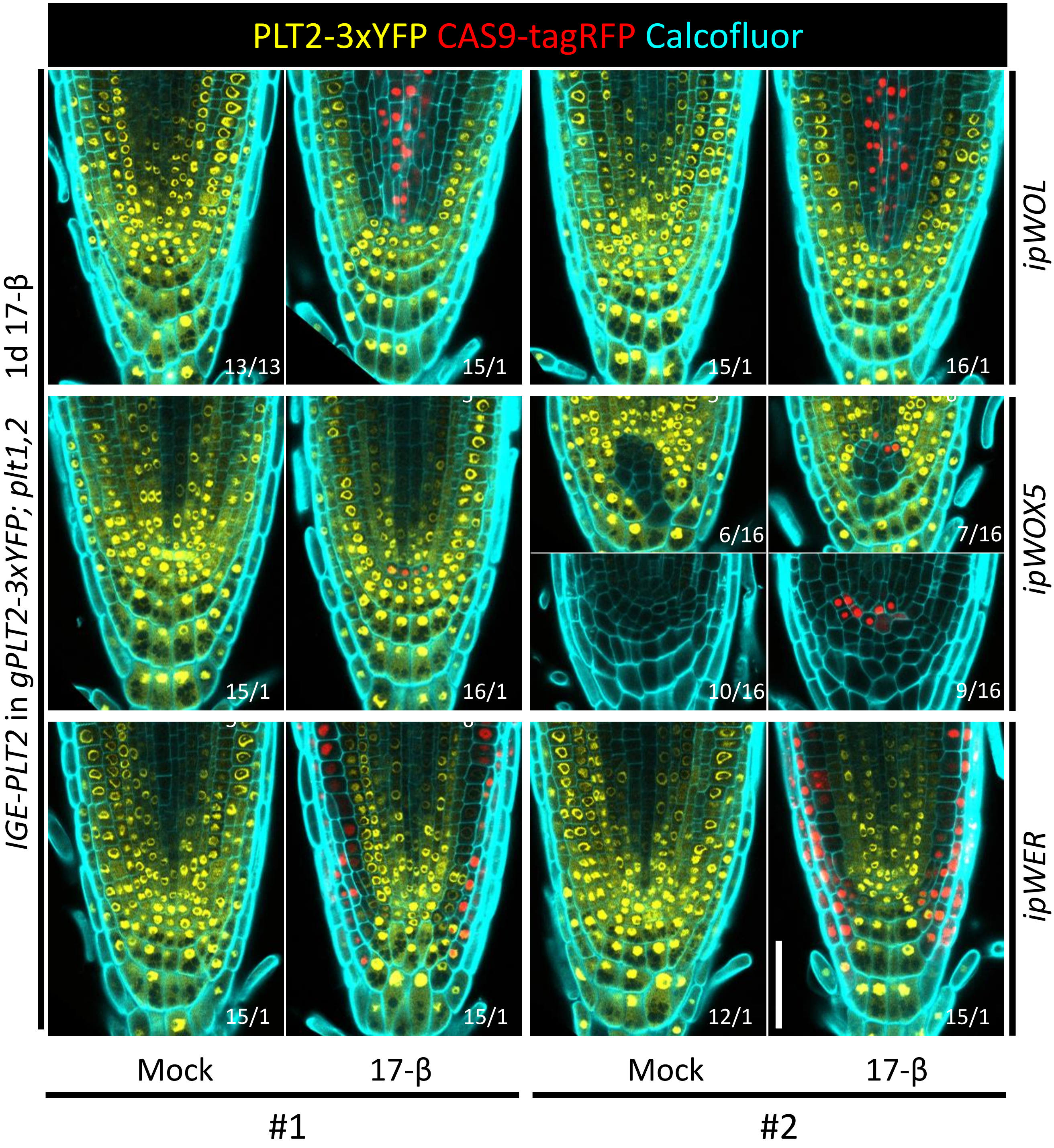


**Supplementary Figure 6 The capacity of conditional genome editing by IGE system is inherited.**

For each construct, two independent transgenic T2 lines were randomly selected and checked. Representative images are shown. Note that the second *ipWOX5>>Cas9p-tagRFP-PLT2* line was leaky: roots displayed a similar phenotype with/without induction. Cell walls are marked by calcofluor. Numbers represent the frequency of the observed phenotype in analyzed T2 samples. Scale bar, 50 µm.

### Supplementary Figure 7

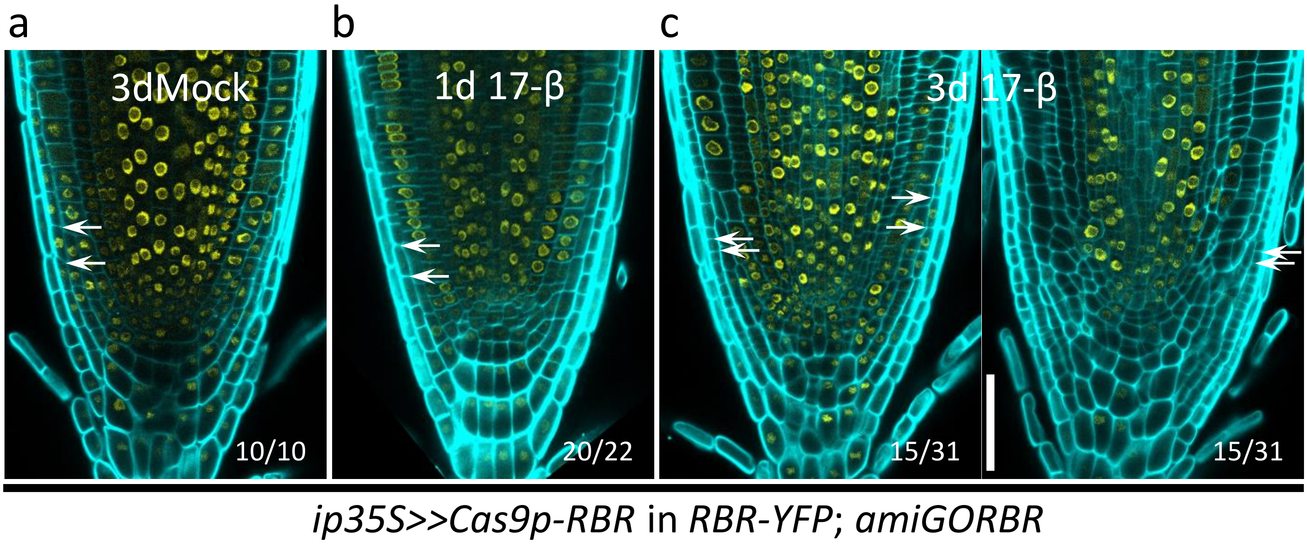


**Supplementary Figure 7 RBR functions cell-autonomously in the RM.**

**(a)** A three-day mock treatment of *ip35S>>Cas9p-RBR* in *RBR-YFP; amiGORBR*. **(b)** A one-day induction caused a reduced RBR-YFP signal mainly in the root cap region without an obvious phenotype. **(c)** Inducing *RBR* editing with *ip35S* typically led to LRC overproliferation (white arrows) without affecting the YFP signal in other domains after a 3-day induction. In some cases, both wild type cells and RBR-knockout cells were seen on the same root (left in **c**). Cell walls are visualized by calcofluor. Numbers indicate the frequency of the observed phenotype in independent T1 samples. Scale bar, 50 μm.

### Supplementary Figure 8

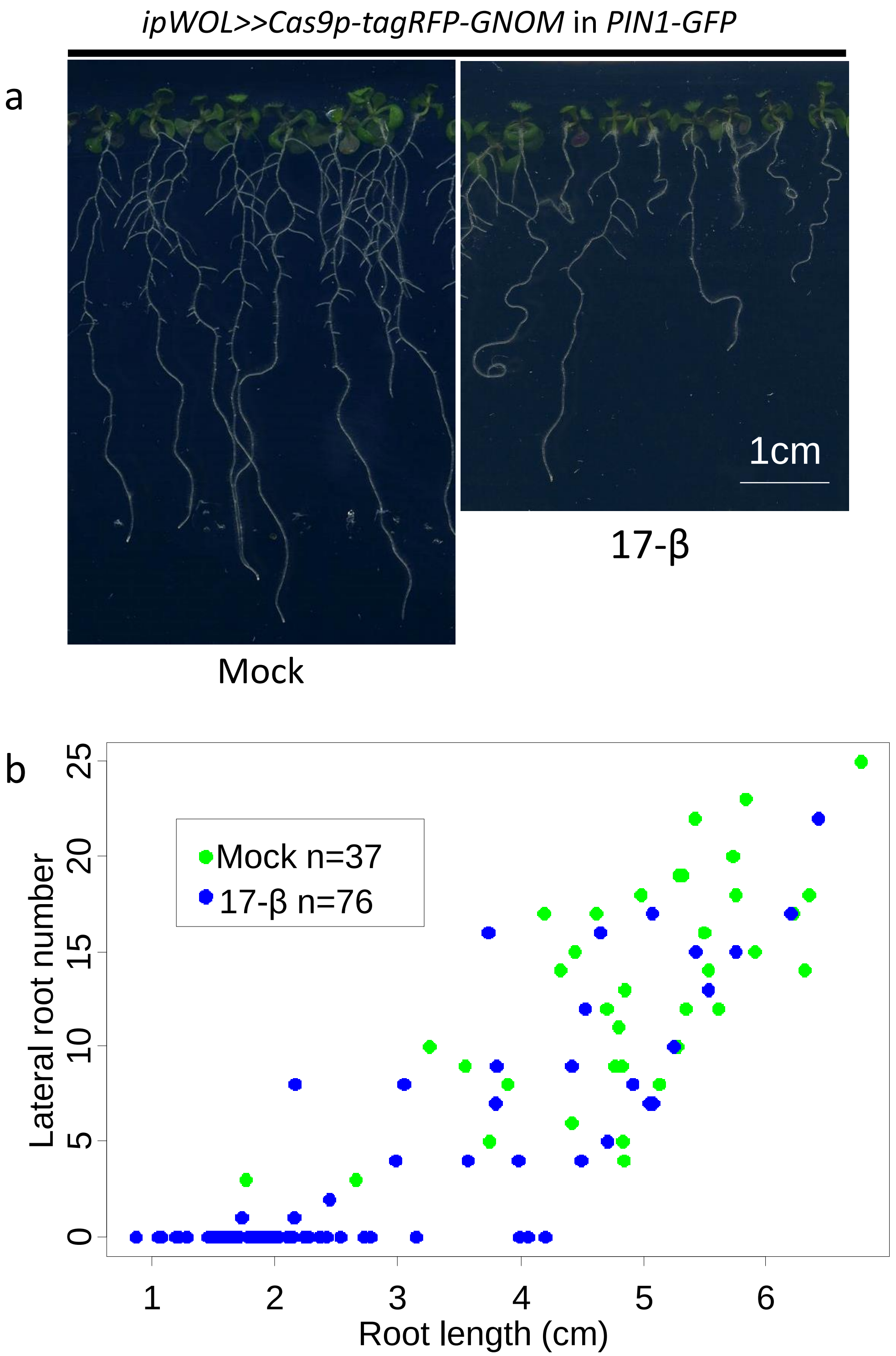


**Supplementary Figure 8 Post-embryonically inducing *GNOM* editing recapitulates the phenotypes of the *gnom* mutant.**

**(a)** Plants with *ipWOL>>Cas9p-tagRFP-GNOM*; *PIN1-GFP* ten days after germination on mock or 17-β plates. Inducing *GNOM* editing led to shorter roots, agravitropic growth and decreased lateral root (LR) numbers. Adventitious roots from the hypocotyl were frequently found, but these roots were not counted in LR quantification. For each independent root, LR number and root length is quantified in **(b)**. Scale bar, 1 cm.

### Supplementary Figure 9

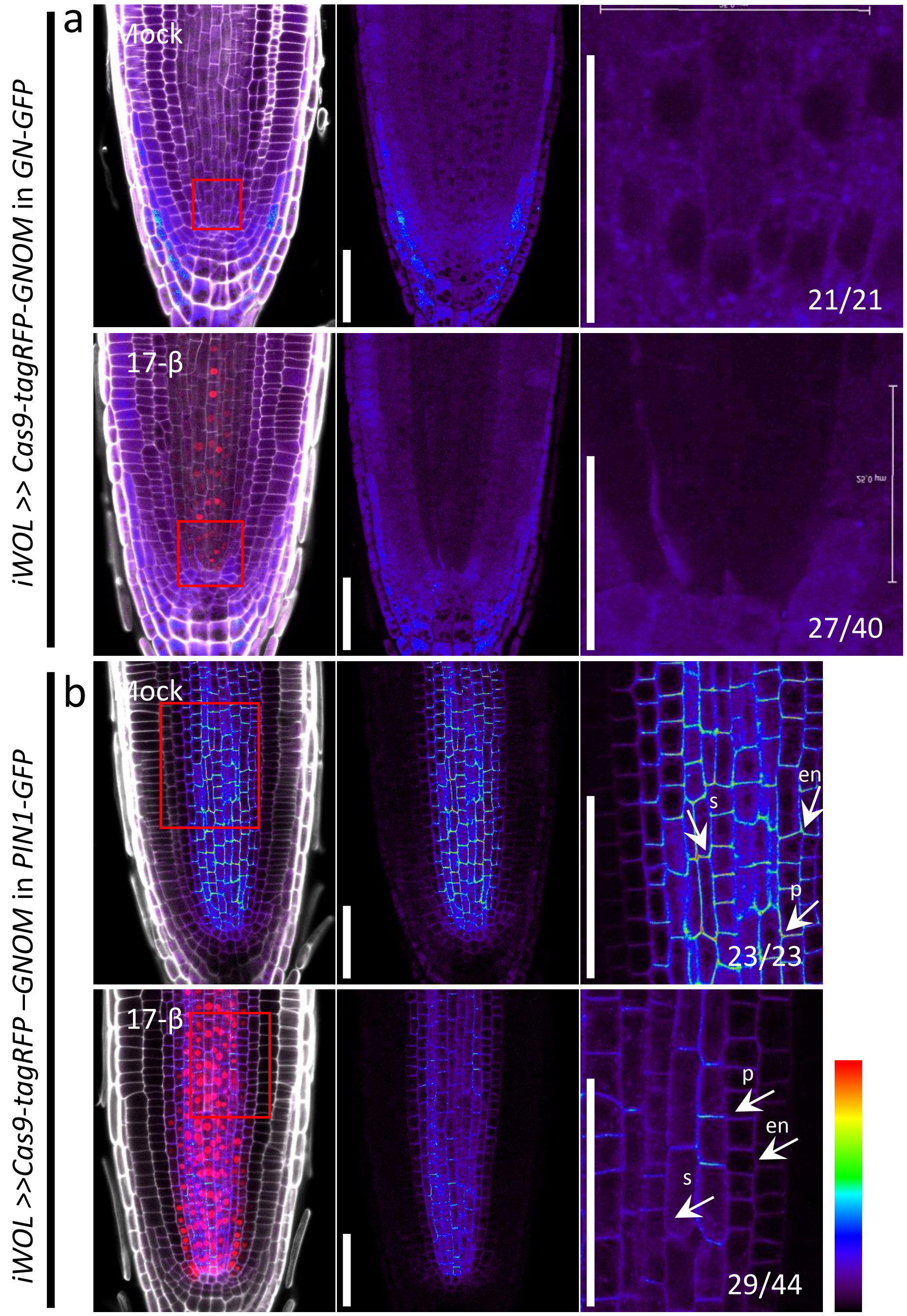


**Supplementary Figure 9 GNOM is required for PIN1 polarity and expression.**

**(a)** *GNOM* expression disappeared from the vasculature after a 6-day induction of IGE targeting *GNOM*. Due to the weak GFP signal, only roots showing a clear loss of GFP signal were included in quantification. **(b)** A three-day induction of *ipWOL>> Cas9p-tagRFP-GNOM*; *PIN1-GFP* resulted in loss of polarity and decreased expression of PIN1-GFP in the endodermis (en), pericycle (p) and stele (s) (white arrows). Right panels are magnified images of the regions marked with a red box in the left panels. Cell walls are marked by calcofluor. Numbers indicate the frequency of the observed phenotype in independent T1 samples analyzed. Scale bar in right panels of **a**, 25 μm; others, 50 μm.

### Supplementary Figure 10

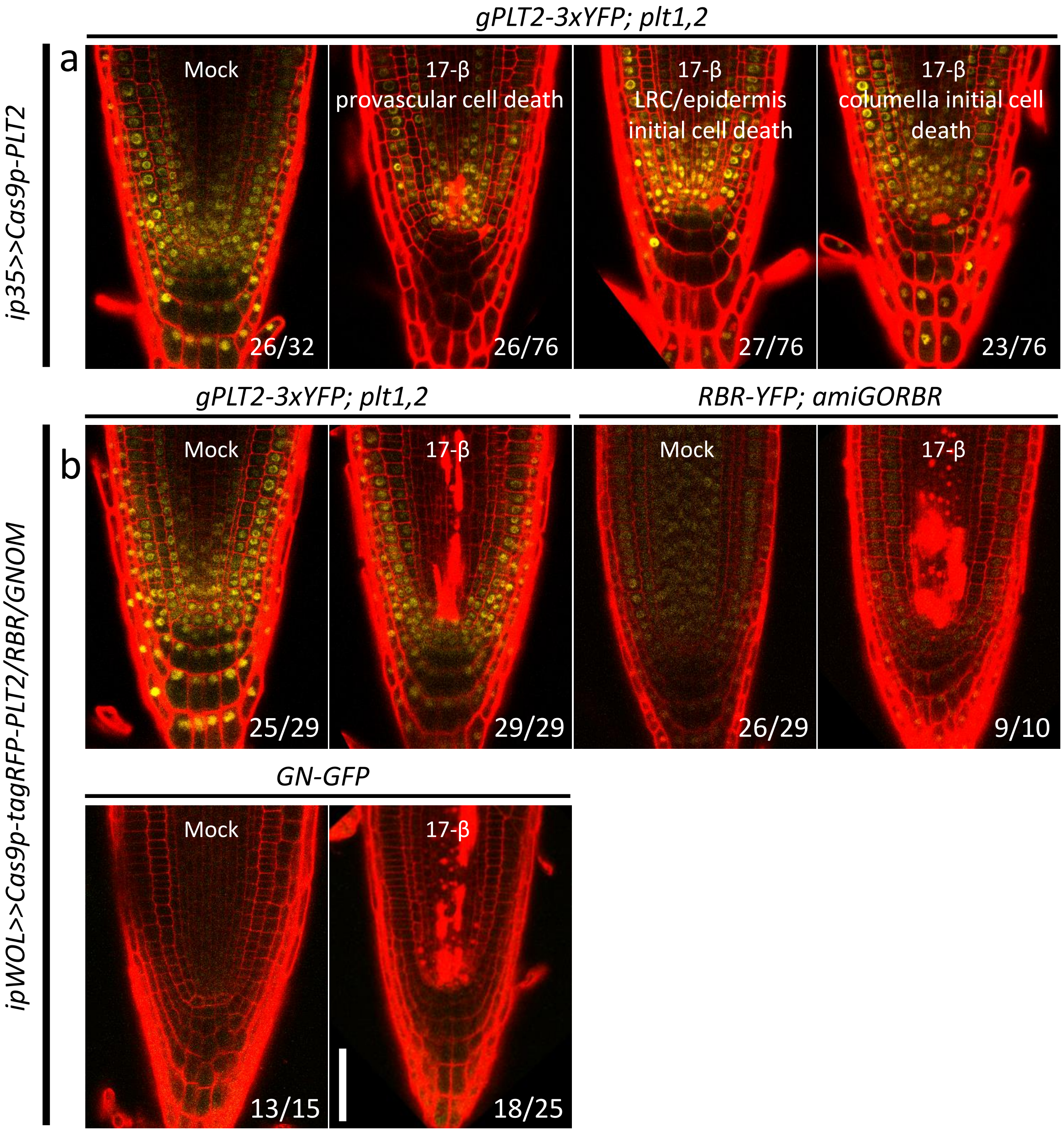


**Supplementary Figure 10 Cas9p-mediated genome editing in proximal stem cells induces cell death.**

**(a)** Stem cell death surrounding the QC was observed after one day of *ip35S>>Cas9p-PLT2* induction. Based on cell types, the cell death response is classified into three categories: provascular cell death, LRC/epidermis initial cell death and columella initial cell death. Samples were counted twice if they had cell death in different domains. **(b)** Cell death of provascular cells and early descendants was induced after one day of induction of *ipWOL>>Cas9p-tagRFP-PLT2/RBR/GNOM*. Cell walls are highlighted by propidium iodide (PI). Under PI detection settings, Cas9p-tagRFP is also visible. Numbers indicate the frequency of the observed phenotype in independent T1 samples analyzed. Scale bars, 50 µm.

### Supplementary Figure 11

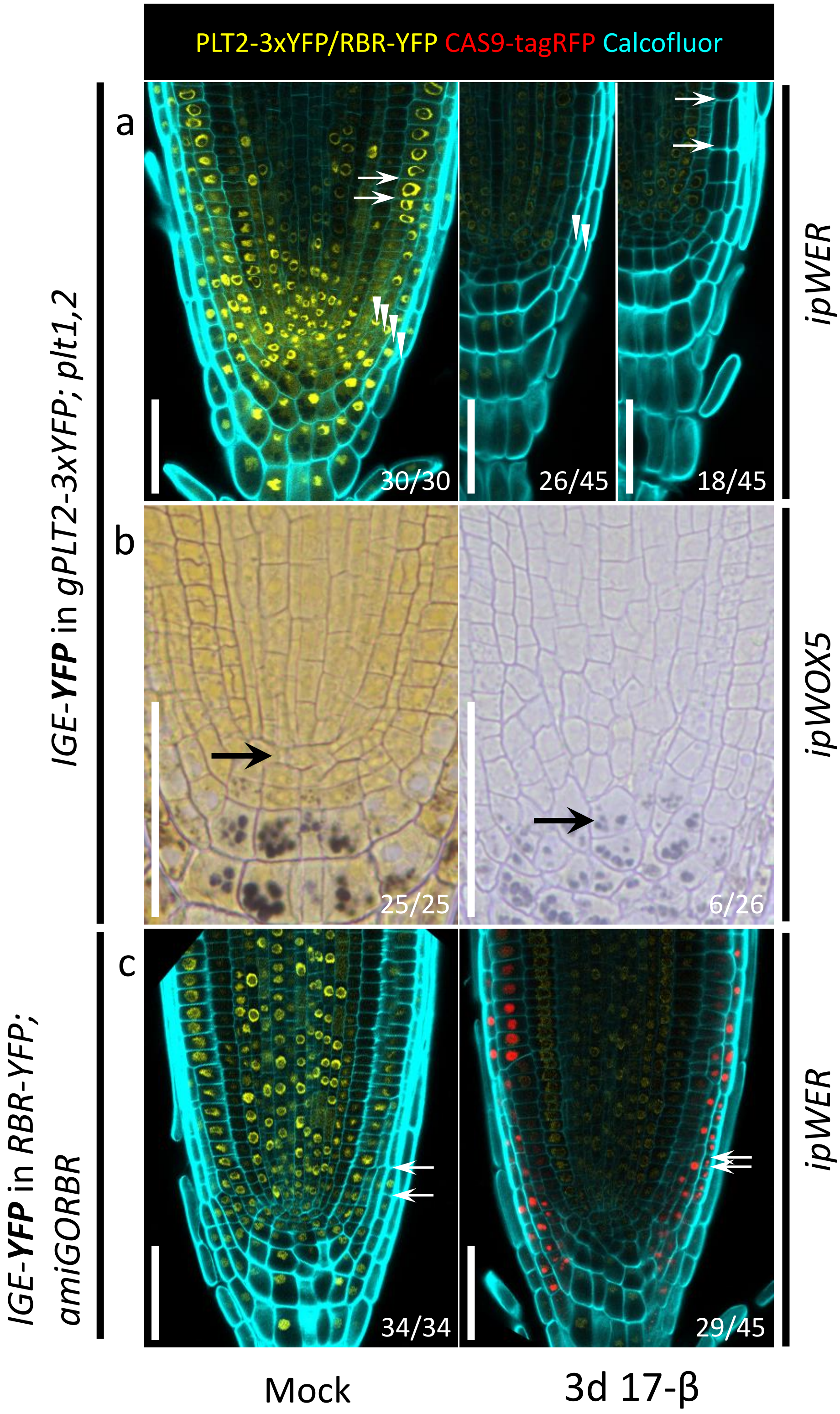


**Supplementary Figure 11 A single IGE construct targeting a gene encoding a fluorescent reporter has the potential to disrupt different transgene targets.**

**(a)** Editing *YFP* instead of *PLT2* in the *ipWER* expression region caused changes similar to direct *PLT2* editing. The RM had fewer LRC layers (white arrowheads), as well as premature expansion of epidermal cells and a broad, faint YFP signal. The Cas9p-tagRFP signal is frequently invisible. **(b)** Editing *YFP* led to QC (black arrow) differentiation at a lower frequency. **(c)** Targeting the *YFP* of RBR-YFP in the LRC led to LRC overproliferation, similar to editing RBR. However, the YFP signal outside *ipWER* expression region was also hampered by an unknown mechanism, unlike when editing *RBR*. White arrows mark the neighboring cell walls in **a** and **c**. The same construct was used in **a** and **c**. Cell walls are highlighted by calcofluor. Numbers indicate the frequency of the observed phenotype in independent T1 samples analyzed. Scale bars, 50 μm.

### Supplementary Figure 12

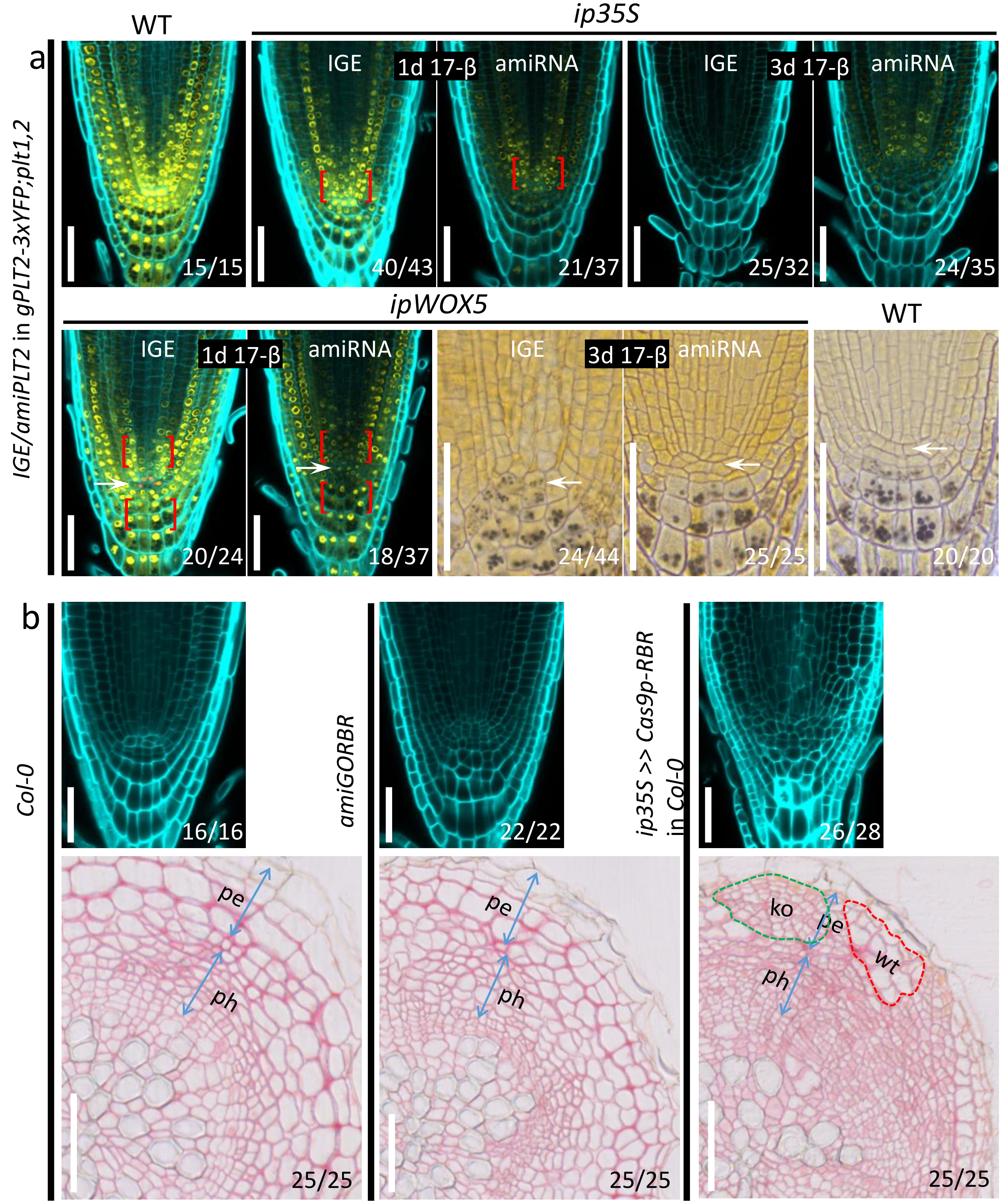


**Supplementary Figure 12 Comparison of IGE system with inducible amiRNA.**

**(a)** IGE-PLT2 displays more specific and stronger *PLT2-YFP* downregulation than amiPLT2. After a one-day induction, *ip35S>>amiPLT2-1*; *gPLT2-3xYFP*;*plt1,2* and *ipWOX5>>amiPLT2-1*; *gPLT2-3xYFP*;*plt1,2* showed a broader reduction of the YFP signal, particularly in the bracketed regions where no inducible promoter activity was found. Conversely, induced *PLT2* editing caused very local loss of the YFP signal. After a three-day induction, the YFP signal is still visible in most of *ip35S>>amiPLT2-1*; *gPLT2-3xYFP; plt1,2* transformants but not in *ip35S>>Cas9p-PLT2*; *gPLT2-3xYFP*;*plt1,2* transformants. There was no QC differentiation in *ipWOX5>>amiPLT2-1*; *gPLT2-3xYFP*; *plt1,2* roots. WT here means 7-day old *gPLT2-3xYFP; plt1,2*. White arrows mark the QC. **(b)** Comparison of the RM and root secondary growth of *Col-0, 35S:amiGORBR* and *ip35S>>Cas9p-RBR*. Inducing *RBR* editing (germination and six days of growth on 17-β plates) resulted in more excessive cell divisions in the LRC than was seen in *amiGORBR* roots (germination and six days of growth on 17-β free plates). Furthermore, RBR editing caused cell overproliferation in secondary tissues such as phloem (ph) cells and the periderm (pe), which was not observed in *amiGORBR* roots. The knockout (ko) sectors (green dotted line) were frequently accompanied by WT sectors (red dotted line), which can be regarded as an internal control. Cell walls are marked by calcofluor. Numbers indicate the frequency of observed phenotype in independent samples analyzed. Scale bars, 50 µm.

**Supplementary Table 1 Quantification of fully differentiated RM after 10 days induction.**

| 1st BOX | 2nd BOX | 3rd BOX | Differentiated RM after 10d 17-β induction. Two repeats | |
|---|---|---|---|---|
| p1R4-35S-XVE | p221z-CAS9p | p2R3z-PLT2-AtU3b-sgRNA1 | 31/47 (66.0 %) | 25/41 (61.0 %) |
| | | p2R3z-PLT2-AtU3d-sgRNA1 | 17/32 (53,1 %) | 20/48 (41,7 %) |
| | | p2R3z-PLT2-AtU6-1-sgRNA1 | 0/29 (0.0 %) | 0/43 (0.0 %) |
| | | p2R3z-PLT2-AtU6-29-sgRNA1 | 15/23 (65.2 %) | 22/34 (64.7 %) |
| | | p2R3z-PLT2-AtU3b-tRNA-sgRNA1 | 20/34 (58.8 %) | 25/31 (80.6 %) |
| | | p2R3z-PLT2-AtU3b-sgRNA1+AtU3d-sgRNA2+AtU6-1-sgRNA3+AtU6-29-sgRNA4 | 17/32 (53.1 %) | 25/35 (71.4 %) |
| | p221z-CAS9p-taqRFP | p2R3z-PLT2-AtU3b-sgRNA1 | 21/32 (65.6 %) | 23/39 (59.0 %) |
| | p221z-dCAS9p | p2R3z-PLT2-AtU3b-sgRNA1+AtU3d-sgRNA2+AtU6-1-sgRNA3+AtU6-29-sgRNA4 | 0/32 (0.0 %) | 0/41 (0.0 %) |
| | p221z-AtMIR390-PLT2-1 | p2R3z-nosT2 | 0/29 (0.0 %) | 0/32 (0.0 %) |
| | p221z-AtMIR390-PLT2-2 | p2R3z-nosT2 | 0/24 (0.0 %) | 0/37 (0.0 %) |

**Supplementary Table 2 Primer list in this study.**

| Primer name | sequence(5'-3') | purpose |
|---|---|---|
| attB1-CAS9p-T35s-F | GGGGACAAGTTTGTACAAAAAAGCAGGCTCGATGGCTCCTAAGAAGAAGCG | For cloning CAS9p with T35s terminator into 2nd BOX |
| attB2-CAS9p-T35s-R | GGGGACCACTTTGTACAAGAAAGCTGGGTGGTCACTGGATTTTGGTTTTAGG | |
| attB2-ccdB-F | GGGGACAGCTTTCTTGTACAAAGTGGAACTCGAGAGACCTCTGAAGTGG | clone Bsa I-ccdB-Bsa I into 3 box |
| attB3-ccdB-R | GGGGACAACTTTGTATAATAAAGTTGAACCGCGAGACCCACGCTCAC | |
| PLT2-TG1-gRT#+ | TGTGAAGAGTGAATGTGAGGGTTTTAGAGCTAGAAAT | clone 4 sgRNA expression cassettes targeting PLT2 |
| PLT2-TG1-AtU3bT#- | CCTCACATTCACTCTTCACATGACCAATGTTGCTCC | |
| PLT2-TG2-gRT#+ | ATAAGGTACGAGGTTGTGATGTTTTAGAGCTAGAAAT | |
| PLT2-TG2-AtU3dT#- | ATCACAACCTCGTACCTTATTGACCAATGGTGCTTTG | |
| PLT2-TG3-gRT#+ | TTAGATAACTAACTACGAGAGTTTTAGAGCTAGAAAT | |
| PLT2-TG3-AtU6-1T#- | TCTCGTAGTTAGTTATCTAACAATCACTACTTCGTCT | |
| PLT2-TG4-gRT#+ | CATCAATATGGTGCAGCGAGGTTTTAGAGCTAGAAAT | |
| PLT2-TG4-AtU6-29T#- | CTCGCTGCACCATATTGATGCAATCTCTTAGTCGACT | |
| dCas9p-D10A-F | TACTCCATCGGCCTCgcgATCGGCACCAACAGC | dCas9 cloning |
| dCas9p-H840A-R | GACTGAGGAACAATcgcGTCGACGTCGTAGT | |
| attB1-gPLT2-F | GGGGACAAGTTTGTACAAAAAAGCAGGCTCGATGAATTCTAACAACTGGCTC | PCR detection of PLT2 deletion from genome, and subsequent cloning into pDONR221z for sequencing |
| attB2-gPLT2-R1 | GGGGACCACTTTGTACAAGAAAGCTGGGTGGAATCATGATACTGAGAGAT | |
| attB2-gPLT2-R2 | GGGGACCACTTTGTACAAGAAAGCTGGGTGGAGCTTGACCCAATACCAAT | |
| attB2-gPLT2-R3 | GGGGACCACTTTGTACAAGAAAGCTGGGTGGATCCTTGAGCAGACTCTCC | |
| amiPLT2-1-F | TGTATGATGATCCCCCGATTTGCTGATGATGATCACATTCGTTATCTATTTTTTCAGCAAATCGTGGGATCATCA | amiPLT2-1 cloning |
| amiPLT2-1-R | AATGTGATGATCCCACGATTTGCTGAAAAAATAGATAACGAATGTGATCATCATCAGCAAATCGGGGGATCATCA | |
| amiPLT2-2-F | TGTATGATCGGTGTGATGATCCCCGATGATGATCACATTCGTTATCTATTTTTTCGGGGATCATAACACCGATCA | amiPLT2-2 cloning |
| amiPLT2-2-R | AATGTGATCGGTGTTATGATCCCCGAAAAAATAGATAACGAATGTGATCATCATCGGGGATCATCACACCGATCA | |
| PLT2-TG1-AtU3dT#- | CCTCACATTCACTCTTCACATGACCAATGGTGCTTTG | sgRNA promoter comparison |
| PLT2-TG1-AtU6-1T#- | CCTCACATTCACTCTTCACACAATCACTACTTCGTCT | |
| PLT2-TG1-AtU6-29T#- | CCTCACATTCACTCTTCACACAATCTCTTAGTCGACT | |
| YFP-gRT | CCCATCCTGGTCGAGCTGGAGTTTTAGAGCTAGAAAT | YFP targeting |
| AtU3b-YFP | TCCAGCTCGACCAGGATGGGTGACCAATGTTGCTCC | |
| RBR-TG1-gRT#+ | TCAGCAAGCATGTCTAACATGTTTTAGAGCTAGAAAT | For cloning 4 sgRNA expression cassettes targeting RBR |
| RBR-TG1-AtU3bT# | ATGTTAGACATGCTTGCTGATGACCAATGTTGCTCC | |
| RBR-TG2-gRT#+ | GTCAAGGCTGGATCTGTACTGTTTTAGAGCTAGAAAT | |
| RBR-TG2-AtU3dT# | AGTACAGATCCAGCCTTGACTGACCAATGGTGCTTTG | |
| RBR-TG3-gRT#+ | TATCCTCAACTCATCTTCTGGTTTTAGAGCTAGAAAT | |
| RBR-TG3-AtU6-1T# | CAGAAGATGAGTTGAGGATACAATCACTACTTCGTCT | |
| RBR-TG4-gRT#+ | TATGACAGTCCTGAGCCACTGTTTTAGAGCTAGAAAT | |
| RBR-TG4-AtU6-29T# | AGTGGCTCAGGACTGTCATACAATCTCTTAGTCGACT | |
| GNOM-TG1-gRT#+ | ACTACACTTGTCAACAGAGCGTTTTAGAGCTAGAAAT | For cloning 4 sgRNA expression cassettes targeting GNOM |
| GNOM-TG1-AtU3bT# | GCTCTGTTGACAAGTGTAGTTGACCAATGTTGCTCC | |
| GNOM-TG2-gRT#+ | TTGATGGATGATGGACCAGTGTTTTAGAGCTAGAAAT | |
| GNOM-TG2-AtU3dT# | ACTGGTCCATCATCCATCAATGACCAATGGTGCTTTG | |
| GNOM-TG3-gRT#+ | GTGTACTCATCAAGATGGACGTTTTAGAGCTAGAAAT | |
| GNOM-TG3-AtU6-1T# | GTCCATCTTGATGAGTACACCAATCACTACTTCGTCT | |
| GNOM-TG4-gRT#+ | TCAGCTCATCTACAGTCAATGTTTTAGAGCTAGAAAT | |
| GNOM-TG4-AtU6-29T# | ATTGACTGTAGATGAGCTGACAATCTCTTAGTCGACT | |

| | | |
|---|---|---|
| attB2-AtU3b-F | GGGGACAGCTTTCTTGTACAAAGTGGAATTTACTTTAAATTTTTTCTTAT | Generating p2R3z-AtU3b-tRNA-ccdB-gRNA entry clone |
| tRNA-AtU3b-R | ACCACTAGACCACTGGTGCTTTGTTTGACCAATGTTGCTCCCTCAGTGTT | |
| AtU3b-tRNA-F | TAACACTGAGGGAGCAACATTGGTCAAACAAAGCACCAGTGGTCTA | |
| tRNA-R | CCGTGGCAGGGTACTATTCTACCACTAGACCACTGGTGCTTTGTT | |
| tRNA-F | AGAATAGTACCCTGCCACGGTACAGACCCGGGTTCGATTCCCGGCT | |
| ccdB-tRNA-R | TGAATCGGCCACTTCAGAGGTCTCTTGCACCAGCCGGGAATCGAACCCGGG | |
| tRNA-ccdB-F | CCCGGGTTCGATTCCCGGCTGGTGCAAGAGACCTCTGAAGTGGCCGATTCA | |
| ccdB-sgRNA-R | AACTTGCTATTTCTAGCTCTAAAACCGAGACCCACGCTCACCCGCCGCGC | |
| ccdB-sgRNA-F | GCGCGGCGGGTGAGCGTGGGTCTCGGTTTTAGAGCTAGAAATAGCAAGTT | |
| attB3-sgRNA-R | GGGGACAACTTTGTATAATAAAGTTGAAAAAAAAAAGCACCGACTCGGTGCCA | |
| BSAI-PLT2-TG1-F | TGCATGTGAAGAGTGAATGTGAGG | For cloning PLT2 target 1 into 2R3z-AtU3b-tRNA-CCD-gRNA entry clone |
| BSAI-PLT2-TG1-R | AAACCCTCACATTCACTCTTCACA | |
| CAS9-RFP-F | CGTATCGACCTTTCCCAGCTTGGTGGTGATATGAGCGAGCTGATTAAGGA | For making p221z-Cas9-tagRFP entry clone |
| NLS-RFP-R | TCCGGCCTTTTTGGTGGCAGCAGGACGCTTCTTGTGCCCCAGTTTGCTAG | |

Underlined sequences indicate Gateway adaptors. Sequence in red represent the target sequence in the gene.

**Supplementary Table 3 Constructs list in this study.**

| expression vector name | 1st BOX | 2nd BOX | 3rd BOX | destination vector |
|---|---|---|---|---|
| 35S:XVE>>CAS9p-PLT2-AtU3b-sgRNA1 | p1R4-35S-XVE | p221z-CAS9p-T35S | p2R3z-PLT2-AtU3b-sgRNA1 | pBm43GW |
| 35S:XVE>>CAS9p-PLT2-AtU3d-sgRNA1 | p1R4-35S-XVE | p221z-CAS9p-T35S | p2R3z-PLT2-AtU3d-sgRNA1 | pBm43GW |
| 35S:XVE>>CAS9p-PLT2-AtU6-1-sgRNA1 | p1R4-35S-XVE | p221z-CAS9p-T35S | p2R3z-PLT2-AtU6-1-sgRNA1 | pBm43GW |
| 35S:XVE>>CAS9p-PLT2-AtU6-29-sgRNA1 | p1R4-35S-XVE | p221z-CAS9p-T35S | p2R3z-PLT2-AtU6-29-sgRNA1 | pBm43GW |
| 35S:XVE>>CAS9p-PLT2-AtU3b-tRNA-sgRNA1 | p1R4-35S-XVE | p221z-CAS9p-T35S | p2R3z-PLT2-AtU3b-tRNA-sgRNA1 | pFRm43GW |
| 35S:XVE>>CAS9p-PLT2-sgRNA1-4 | p1R4-35S-XVE | p221z-CAS9p-T35S | p2R3z-PLT2-AtU3b-sgRNA1+AtU3d-sgRNA2+AtU6-1-sgRNA3+AtU6-29-sgRNA4 | pBm43GW |
| 35S:XVE>>dCAS9p-PLT2-sgRNA1-4 | p1R4-35S-XVE | p221z-dCAS9p-T35S | p2R3z-PLT2-AtU3b-sgRNA1+AtU3d-sgRNA2+AtU6-1-sgRNA3+AtU6-29-sgRNA4 | pBm43GW |
| 35S:XVE>>CAS9p-tagRFP-PLT2-AtU3b-sgRNA1 | p1R4-35S-XVE | p221z-CAS9p-tagRFP-T35S | p2R3z-PLT2-AtU3b-sgRNA1 | pBm43GW |
| 35S:XVE>>AtMIR390-PLT2-1-nosT2 | p1R4-35S-XVE | p221z-AtMIR390-PLT2-1 | nosT2 | pFRm43GW |
| 35S:XVE>>AtMIR390-PLT2-2-nosT2 | p1R4-35S-XVE | p221z-AtMIR390-PLT2-2 | nosT2 | pFRm43GW |
| pWOX5:XVE>>AtMIR390-PLT2-1-nosT2 | p1R4-pWOX5:XVE | p221z-AtMIR390-PLT2-1 | nosT2 | pFRm43GW |
| pWER:XVE>>CAS9p-tagRFP-PLT2-sgRNA1-4 | p1R4-pWER:XVE | p221z-CAS9p-tagRFP-T35S | p2R3z-PLT2-AtU3b-sgRNA1+AtU3d-sgRNA2+AtU6-1-sgRNA3+AtU6-29-sgRNA4 | pBm43GW |
| pWOX5:XVE>>CAS9p-tagRFP-PLT2-sgRNA1-4 | p1R4--pWOX5:XVE | p221z-CAS9p-tagRFP-T35S | p2R3z-PLT2-AtU3b-sgRNA1+AtU3d-sgRNA2+AtU6-1-sgRNA3+AtU6-29-sgRNA4 | pBm43GW |
| pSCR:XVE>>CAS9p-tagRFP-PLT2-sgRNA1-4 | p1R4--pSCR:XVE | p221z-CAS9p-tagRFP-T35S | p2R3z-PLT2-AtU3b-sgRNA1+AtU3d-sgRNA2+AtU6-1-sgRNA3+AtU6-29-sgRNA4 | pBm43GW |
| pWOL:XVE>>CAS9p-tagRFP-PLT2-sgRNA1-4 | p1R4-pWOL:XVE | p221z-CAS9p-tagRFP-T35S | p2R3z-PLT2-AtU3b-sgRNA1+AtU3d-sgRNA2+AtU6-1-sgRNA3+AtU6-29-sgRNA4 | pBm43GW |
| pWER:XVE>>CAS9p-taRFP-RBR-sRNA1-4 | p1R4-pWER:XVE | p221z-CAS9p-tagRFP-T35S | p2R3z-RBR-AtU3b-sgRNA1+AtU3d-sgRNA2+AtU6-1-sgRNA3+AtU6-29-sgRNA4 | pFRm43GW |
| pWOX5:XVE>>CAS9p-taRFP-RBR-sRNA1-4 | p1R4-pWOX5:XVE | p221z-CAS9p-tagRFP-T35S | p2R3z-RBR-AtU3b-sgRNA1+AtU3d-sgRNA2+AtU6-1-sgRNA3+AtU6-29-sgRNA4 | pFRm43GW |
| pSCR:XVE>>CAS9p-taRFP-RBR-sRNA1-4 | p1R4--pSCR:XVE | p221z-CAS9p-tagRFP-T35S | p2R3z-RBR-AtU3b-sgRNA1+AtU3d-sgRNA2+AtU6-1-sgRNA3+AtU6-29-sgRNA4 | pFRm43GW |
| pWOL:XVE>>CAS9p-taRFP-RBR-sRNA1-4 | p1R4-pWOL:XVE | p221z-CAS9p-tagRFP-T35S | p2R3z-RBR-AtU3b-sgRNA1+AtU3d-sgRNA2+AtU6-1-sgRNA3+AtU6-29-sgRNA4 | pFRm43GW |
| 35S:XVE>>CAS9p-RBR-sgRNA1-4 | p1R4-35S-XVE | p221z-CAS9p-T35S | p2R3z-RBR-AtU3b-sgRNA1+AtU3d-sgRNA2+AtU6-1-sgRNA3+AtU6-29-sgRNA4 | pFRm43GW |
| pWER:XVE>>CAS9p-tagRFP-AtU3b-YFP-sgRNA | p1R4-pWER:XVE | p221z-CAS9p-tagRFP-T35S | 2R3z-VEN-AtU3b-sgRNA | pFRm43GW |
| pWOX5:XVE>>CAS9p-tagRFP-AtU3b-YFP-sgRNA | p1R4--pWOX5:XVE | p221z-CAS9p-tagRFP-T35S | 2R3z-VEN-AtU3b-sgRNA | pFRm43GW |
| pWOL:XVE>>CAS9p-tagRFP-GNOM-sgRNA1-4 | p1R4-pWOL:XVE | p221z-CAS9p-tagRFP-T35S | p2R3z-GNOM-AtU3b-sgRNA1+AtU3d-sgRNA2+AtU6-1-sgRNA3+AtU6-29-sgRNA4 | pFRm43GW |
